# Functional analysis of the *Nematostella* Wnt/β-catenin destruction complex provides insight into the evolution of a critical regulatory module in a major metazoan signal transduction pathway

**DOI:** 10.1101/2024.10.27.620500

**Authors:** Hongyan Sun, Brian M. Walters, Radim Zidek, Mark Q. Martindale, Athula H. Wikramanayake

## Abstract

The genesis of signaling pathways likely drove metazoan evolution, but the origins of these pathways and how they acquired signaling activity is poorly understood. Here, we studied functional evolution of the Wnt/β-catenin (cWnt) pathway destruction-complex (DC) that regulates β-catenin signaling. Bilaterian DC function requires β-catenin binding to Axin and APC proteins, and Axin-APC heterodimerization. However, bioinformatic analyses predicted that Axin and APC-like homologs in early-branching non-bilaterians lack important previously defined bilaterian β-catenin binding domains, questioning if they have functional cWnt-DCs. We demonstrate that both Axin and APC proteins in the cnidarian *Nematostella vectensis* (a representative of the sister taxon to bilaterians) can regulate cWnt signaling indicating an active cWnt-DC. Using *in vitro* analyses, we show that NvAxin binds Nvβ-catenin weakly despite lacking the conserved bilaterian Axin β-catenin-binding motif (βcatBM). Using AlphaFold3, we identified two predicted βcatBM-like sequences in NvAxin, one within the Axin-RGS domain and another towards the C-terminus. Similar analysis of placozoan, poriferan, and ctenophore Axin identified single βcatBM-like sequences located within Axin-RGS. We show that ctenophore Axin and β-catenin do not interact and changing a conserved leucine on NvAxin-βcatBM-like motifs to resemble the ctenophore sequence abolished NvAxin-Nvβ-catenin interactions. We propose that an ancestral Axin-RGS sequence acquired low-affinity β-catenin binding early in metazoan evolution, followed by motif duplication in the cnidarian-bilaterian ancestor. In bilaterians, the duplicated βcatBM evolved higher-affinity for β-catenin, while the ancestral sequence was lost. Our results demonstrate how phylogenetic insights, AI tools and functional assays can be used to reconstruct the evolution of complex signaling pathways.

## Introduction

The early development and diversification of multicellular animals (metazoans) is regulated by a small set of conserved signal transduction pathways (Gerhart 1999; Pires-daSilva and Sommer 2003; Richards and Degnan 2009; Babonis and Martindale 2017). It has been hypothesized that the evolution of these signaling pathways was critical for the evolution and radiation of the metazoan last common ancestor, or “urmetazoan”, over 600 million years ago (Pires-daSilva and Sommer 2003; Rokas 2008; Richards and Degnan 2009; Babonis & Martindale, 2017; Richter and King 2013; Ros-Rocher et al. 2021). Despite the critical roles of these pathways in metazoan development and evolution, relatively little is known about their origins or their functional activity in early emerging taxa (Pires-daSilva and Sommer 2003; Babonis & Martindale, 2017; Richards and Degnan 2009). One major constraint to advancing this research has been the difficulty in the care and handling of embryos of early emerging (non-bilaterian) model systems to experimentally manipulate signal transduction pathways. However, the advancement of methodology to molecularly manipulate development in cnidarians (e.g., sea anemones, corals, “jelly fish”), the closest outgroup to the Bilateria (Fig. 1A), and foundational knowledge of Wnt/β-catenin (cWnt) pathway function in these animals and has established cnidarians as one of the best non-bilaterian experimental systems to functionally understand the direction of evolutionary change in the cWnt signaling pathway (Wikramanayake et al. 2003; Lee et al. 2007; Layden et al. 2016; Lebedeva et al. 2021; Houliston et al. 2022; Holzem et al. 2024).

**Figure 1.**
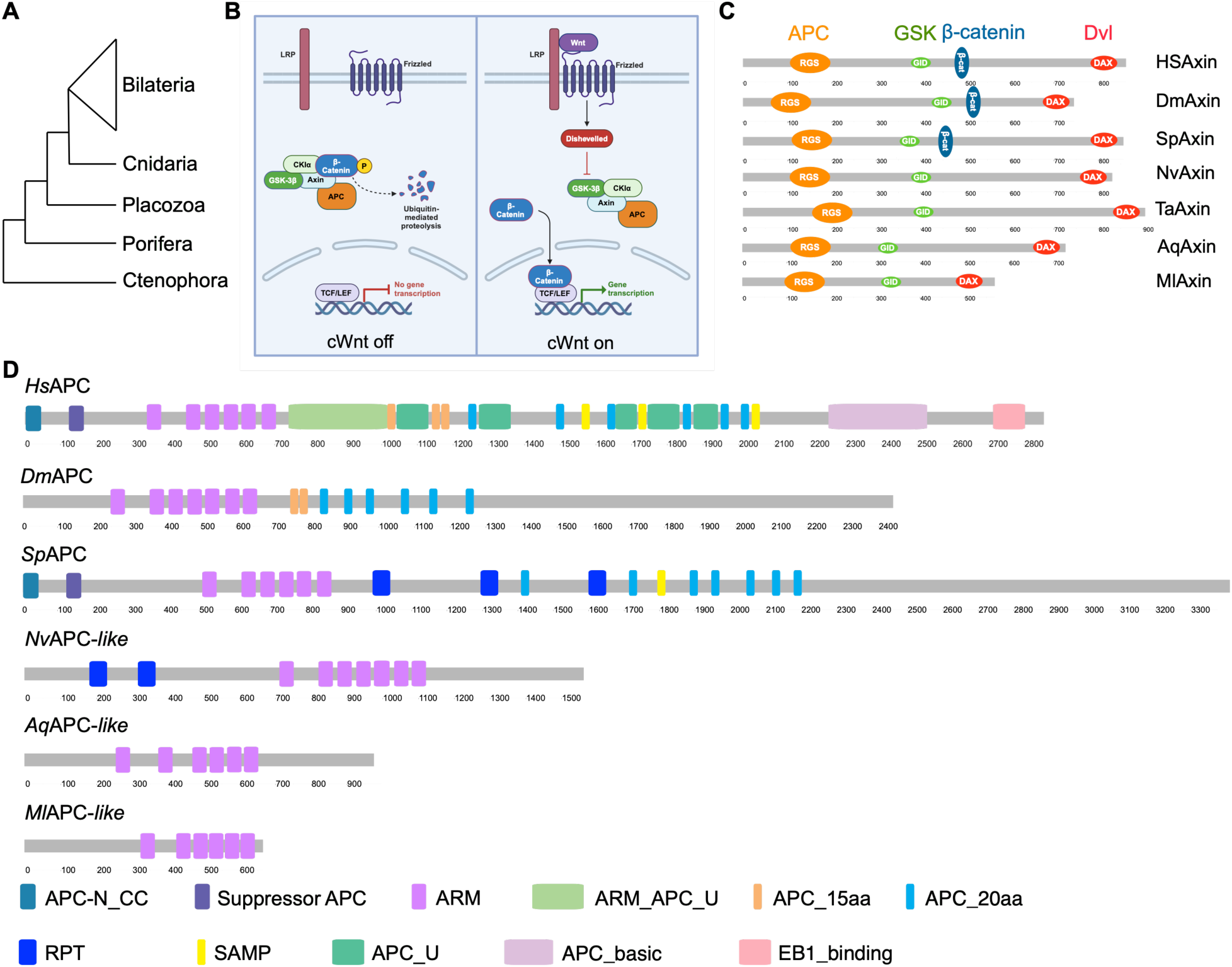
A phylogenetic overview perspective of the evolution of changes in protein-protein interaction domains in the Wnt/ β-catenin pathway destruction complex scaffolding proteins Axin and APC. (A) A simplified phylogenetic tree showing the evolutionary relationships between Bilateria and non-Bilateria taxa. (B) Simplified representation of the cWnt pathway in its “off “and “on” states, showing the activity of the destruction complex scaffolded by the Axin and APC proteins. (C) Schematic diagram comparing the main protein binding domains of Axin in bilaterians and non-bilaterians. (D) Schematic diagram of APC protein domain structures in bilaterians and non-bilaterians. *Hs*, *Homo sapiens*; *Dm*, *Drosophila melanogaster*; *Sp*, *Strongylocentrotus purpuratus*; *Nv*, *Nematostella vectensis*; *Ta*, *Trichoplax adhaerens*; *Aq*, *Amphimedon queenslandica*; *Ml*, *Mnemiopsis leidyi*. The numbers under the protein structures indicate the amino acid position. The color codes for the domains are indicated within or below the protein structures.

The cWnt signaling pathway (Fig. 1B) is a broadly deployed signaling pathway that is restricted to the Metazoa, and cWnt signaling plays diverse roles during evolution, development and homeostasis (Petersen and Reddien 2009; Richards and Degnan 2009; Loh et al. 2016; Babonis and Martindale 2017; Richter et al. 2018; Holzem et al. 2024). Extensive studies of cWnt pathway function in bilaterian models have led to a relatively advanced level of understanding of the critical regulatory nodes in this pathway, hence making it a particularly useful model to investigate how metazoan signaling pathways evolved and acquired signaling competence. For example, like many of the metazoan signal transduction pathways, the cWnt pathway is modular in its organization, with distinct protein complexes regulating transduction of signals from the cell surface to the nucleus (MacDonald et al. 2009; Stamos and Weis 2013; Gammons and Bienz 2018; Liu et al. 2022). One critical cytoplasmic regulatory node in the cWnt pathway is the β-catenin destruction complex (DC) which is essential for regulating cWnt signaling through the control of cytoplasmic levels of β-catenin, the primary transducer of cWnt signaling in the nucleus (Fig. 1B) (Roberts et al. 2011; Stamos and Weis 2013; Gammons and Bienz 2018). Evolution of the DC was critical for the evolution of cWnt signaling because it likely provided a scaffold for exaptation of β-catenin for a signaling function. Moreover, the DC also may have provided a central regulatory node for context-dependent spatial and temporal control of this pathway during its wide deployment in bilaterians during development and homeostasis (Petersen and Reddien 2009; Valenta et al. 2012; Loh et al. 2016; Liu et al. 2022; Holzem et al. 2024).

Axin is a key scaffolding protein in the bilaterian DC which contains multiple binding sites that mediate its direct association with proteins in the DC including β-catenin, the two serine/threonine protein kinases Casein Kinase 1α (CK-1α) and Glycogen Synthase Kinase 3β (GSK-3β), the phosphoprotein Disheveled (Dvl), and APC, the product of the Adenomatous Polyposis Coli gene (Fig. 1C) (Ikeda et al. 1998; Kishida et al. 1998; Fagotto et al. 1999; Valenta et al. 2012; Stamos and Weis 2013; Pronobis et al. 2015; van Kappel and Maurice 2017; Qiu et al. 2024). A detailed model for negative regulation of β-catenin in the cWnt pathway by the DC has emerged from extensive functional studies done in vertebrate and invertebrate model systems (Roberts et al. 2011; Stamos and Weis 2013; van Kappel and Maurice 2017; Liu et al. 2022; Kremer et al. 2010; Peterson-Nedry et al. 2008). This model proposes that β-catenin is recruited to Axin through a conserved binding motif (βcatBM) and this binding mediates the sequential phosphorylation of β-catenin by CK-1α and GSK-3β leading to its ubiquitylation and proteasome-mediated degradation (Xing et al. 2003; Kimelman and Xu 2006; Roberts et al. 2011; Stamos and Weis 2013; van Kappel and Maurice 2017; Gammons and Bienz 2018; Ranes et al. 2021). The transfer of phosphorylated β-catenin to the proteasome requires APC, the second scaffolding protein in the DC. APC in bilaterians is a relatively large protein with multiple domains that have been shown to mediate its interactions with Axin, β-catenin, and over 100 other proteins (Fig. 1D) (Roberts et al. 2011; Nelson and Näthke 2013; van Kappel and Maurice 2017; Abbott and Näthke 2023). Strikingly, APC has multiple distinct binding sites for β-catenin that have different affinities for the protein (Ha et al. 2004; Kimelman and Xu 2006; Roberts et al. 2011; Pronobis et al. 2017; Ranes et al. 2021). Current models postulate that following β-catenin phosphorylation on Axin, one of the 20 amino acid repeats on APC is phosphorylated and subsequently phosphorylated β-catenin is transferred to this domain on APC prior to being targeted to the proteasome for degradation (Kimelman & Xu, 2006; Roberts et al. 2011; Ranes et al. 2021). Regulation of the DC has been subject to extensive experimental interrogation because of its critical role in regulating cWnt signaling, and the discovery that over 90% of human colorectal cancers have mutations in either APC or other components of the DC that can lead to misregulation of cWnt signaling (Polakis 1995; Kinzler and Vogelstein 1996; Fodde et al. 2001; Schneikert and Behrens 2007; Kohler et al. 2008). Oncogenic human APC mutations are frequently seen in the “mutational cluster region” (MCR) that delete many of the well-characterized β-catenin binding sites and the three SAMP domains on the protein that mediate Axin-APC heterodimerization and are required for DC function (Fig. 1D) (Kinzler and Vogelstein 1996; Fodde et al. 2001; Schneikert and Behrens 2007; Kohler et al. 2008; Roberts et al. 2011; Ranes et al. 2021).

The cWnt pathway is deeply conserved with homologs of components of the basic pathway found in all metazoan phyla (Richards and Degnan 2009; Babonis and Martindale 2017; Holzem et al. 2024), but ongoing genomic sequencing studies in early branching metazoan clades have raised interesting fundamental questions about the origin of signaling competence in the cWnt pathway. One of the first observations that raised the possibility that the non-bilaterian cWnt pathway may not function like the bilaterian cWnt pathway came from a phylogenetic analysis of Wnt pathway components in two early emerging taxa, the poriferan (sponge) *Amphimedon queenslandica* and the cnidarian (sea anemone) *Nematostella vectensis* (Richards and Degnan, 2009; Adamska et al. 2010). This work showed that protein domains known to be important in DC proteins in bilaterians were absent in these two species. For example, the Axin proteins from *A. queenslandica* and *N. vectensis* appeared to lack the highly conserved βcatBM seen in bilaterian Axin that mediates its interaction with β-catenin (Richards and Degnan 2009; Adamska et al. 2010) (Fig. 1C). We used the webtool SMART (Simple Modular Architecture Research Tool: http://smart.embl-heidelberg.de) to confirm that Axin from a ctenophore and a placozoan also lack the βcatBM suggesting its bilaterian origin (Fig. 1C). Additionally, APC proteins in these taxa were considerably shorter than the bilaterian homologs, lacking many of the critical protein-protein interaction domains seen in bilaterians (Richards and Degnan 2009; Adamska et al. 2010) (Fig. 1D), and the corresponding genes have recently been classified as “*APC-like*” (Adamska et al. 2010; Abbott and Näthke 2023) suggested that the Axin and APC-like proteins in these early diverging taxa may not be part of the DC, or that these proteins may be interacting in different ways to regulate DC function. These observations provided a fundamental opportunity to better understand the evolution of the regulation of a major metazoan signaling pathway and to reconstruct the ancestral state of a critical regulatory node in the cnidarian-bilaterian last common ancestor (LCA).

In this study we carried out a detailed functional analysis of the cWnt DC in the cnidarian *Nematostella* to determine how it functions in the absence of critical domains in Axin and APC required for bilaterian DC function. We show that *Nematostella* Axin and APC-like proteins use novel protein-protein interactions to form a cWnt DC suggesting that these proteins may have a level of promiscuity not reported in the bilaterian cWnt DC. Additionally, using *in vitro* and *in vivo* protein-protein interaction assays along with AI-enabled prediction we trace the evolutionary history of the critical βcatBM on the bilaterian Axin protein from pre-bilaterian antecedents. We functionally validate the regulation of a non-bilaterian DC function showing that cWnt signaling can achieve deeply conserved developmental outcomes despite significant differences in the regulation of the DC compared to bilaterians. Our results indicate that the key scaffolding proteins in the cWnt DC in the cnidarian-bilaterian last common ancestor were “promiscuous” which provides insight into the mechanisms leading to the evolvability of the signaling pathways in bilaterians promoting the genesis of novel body plans.

## Results

### The conserved β-catenin binding motif in bilaterian Axin proteins is not detectable in non-bilaterian Axins

Structure-function studies in bilaterians have defined conserved regions on the Axin protein that allow it to bind to APC, GSK-3β, and β-catenin in the DC thereby promoting β-catenin phosphorylation and targeting it for degradation, maintaining the pathway in an inactive state (Ikeda et al. 1998; Kishida et al. 1998; Fagotto et al. 1999; Valenta et al. 2012; Stamos and Weis 2013; Pronobis et al. 2015; van Kappel and Maurice 2017; Qiu et al. 2024) (Fig. 1C; Supplementary Table 1). In addition, Axin proteins have a defined domain that binds to “activated” Disheveled when the cWnt pathway is stimulated by upstream signals and a more poorly defined region that interacts with CK-1α (Liu et al. 2002; MacDonald et al. 2009; Gammons and Bienz 2018; Qiu et al. 2024) (Fig. 1C; Supplementary Table 1). These Axin domains play a critical role in recruiting components of the DC to regulate β-catenin stability (Xing et al. 2003; Mendoza-Topaz et al. 2011; Stamos and Weis 2013; Qiu et al. 2024). A previous bioinformatic analysis showed that the βcatBM is absent in *Amphimedon* (Porifera) and *Nematostella* (Cnidaria) Axin proteins (Adamska et al. 2010). To determine if this motif was also absent in the other two non-bilaterian taxa (Placozoa and Ctenophora), the functional domains of Axin proteins in the four non-bilaterian taxa and some selected bilaterian taxa were annotated using the webtool SMART (Simple Modular Architecture Research Tool: http://smart.embl-heidelberg.de) and the Axin proteins were aligned using Clustal Omega. This analysis showed that while the βcatBM was identified in all bilaterian Axins, we could not identify this motif in Axin proteins from the four non-bilaterian taxa (Supplementary Table 1). These observations indicated that the βcatBM may have appeared in Axin after the divergence of the last common ancestor of Bilateria and Cnidaria, or that the domain had diverged during bilaterian evolution to render it undetectable in non-bilaterian Axin using the bioinformatic tools we used. These observations also raised the possibility that in the absence of a βcatBM non-bilaterian Axin would not be able to directly regulate β-catenin stability in the cWnt DC.

### NvAxin and NvAPC-like regulate cWnt signaling in Nematostella embryos

We have previously shown that a bilaterian Axin protein can disrupt early development in *Nematostella* (Kumburegama et al. 2011). But to the best of our knowledge, a functional role for a non-bilaterian Axin in regulating cWnt signaling has not been examined. To study Axin function in *Nematostella*, we first examined the expression pattern of *NvAxin* in eggs and early embryos using whole mount in situ hybridization (WMISH) (Supplementary Fig. 1). WMISH revealed that *NvAxin* was expressed ubiquitously at the egg, cleavage and blastula stages (Supplementary Fig. 1A-C). However, starting at the early gastrula stage, *NvAxin* displayed an asymmetrical expression pattern with the mRNA mostly restricted to the future oral pole and this expression persisted through the planula stage (Supplementary Fig. 1D-F). This expression pattern suggested a function for the NvAxin protein in regulating early pattern formation in *Nematostella*.

We next asked if disrupting *NvAxin* function affected cWnt signaling during early *Nematostella* development. It is now well established that cWnt signaling regulates cell fates along the animal-vegetal axis in *Nematostella* (Wikramanayake et al. 2003; Lee et al. 2007; Röttinger et al. 2012; Lebedeva et al. 2021; Wijesena et al. 2022). Blocking β-catenin signaling using a dominant-negative form of NvTcf that prevents β-catenin from interacting with cis-regulatory elements of cWnt target genes leads to the disruption of normal gastrulation and downregulation of genes in cell types derived from animal half blastomeres—the endomesoderm and pharynx (Röttinger et al. 2012) (Supplementary Fig. 2). To determine if NvAxin regulates cWnt signaling and the nuclearization of β-catenin in embryos, we co-expressed NvAxin-GFP with Nvβ-catenin-mCherry by mRNA injection into zygotes and examined developing embryos for Nvβ-catenin-mCherry expression at the blastula stage. Our results showed that while control embryos co-injected with *GFP* and *Nvβ-catenin-mCherry* mRNA exhibited nuclearization of Nvβ-catenin-mCherry protein on one side of the embryo, embryos co-expressing either NvAxin-GFP or Axin-GFP from the sea urchin *Strongylocentrotus purpuratus* (SpAxin-GFP) and Nvβ-catenin-mCherry did not display any nuclear localization of Nvβ-catenin-mCherry (Fig. 2A-C). This result indicated that NvAxin overexpression blocks nuclearization of Nvβ-catenin-mCherry, presumably by mediating phosphorylation and destabilization of β-catenin within the DC.

**Figure 2.**
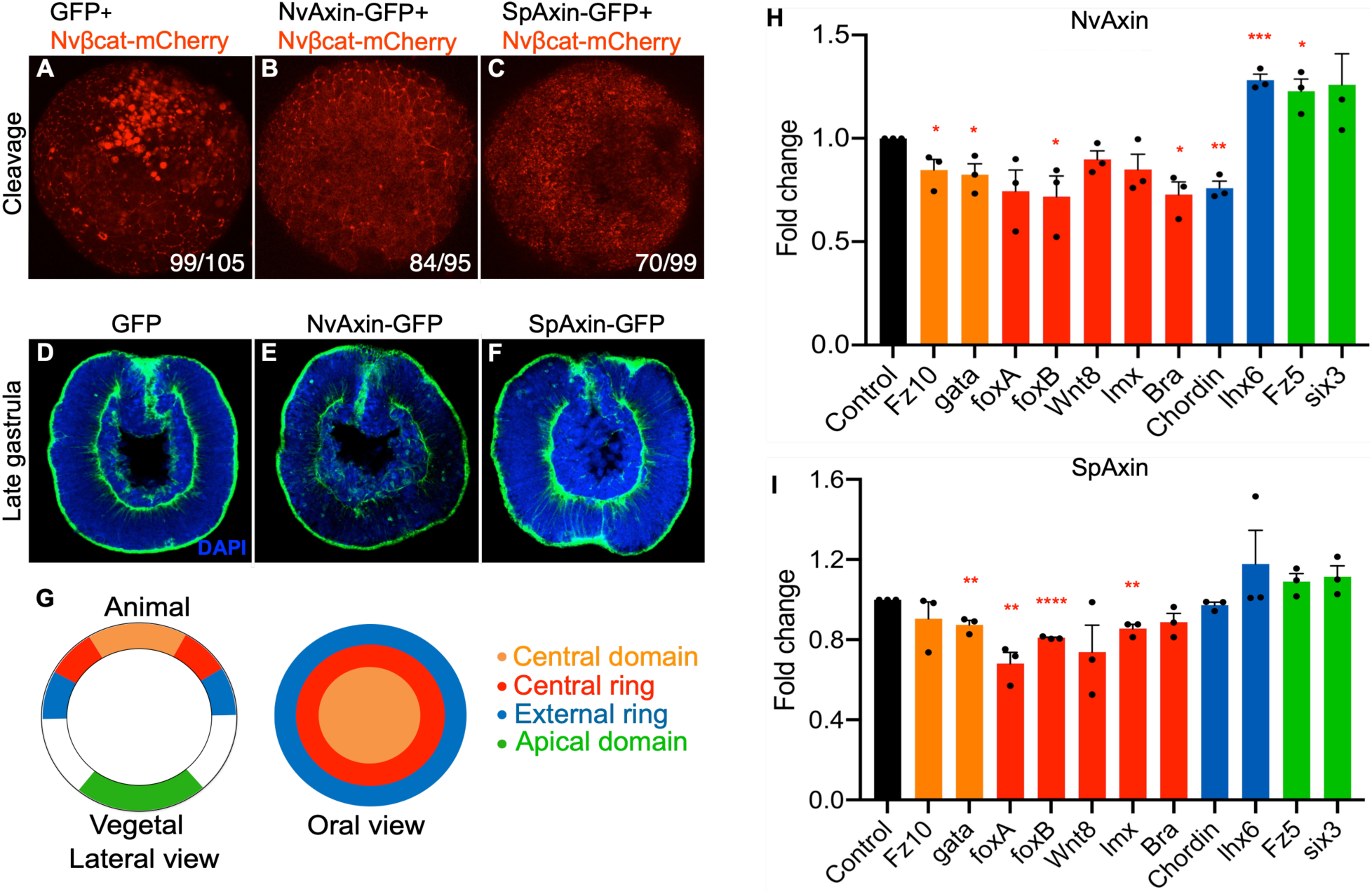
Overexpression of Axin in *Nematostella* embryos disrupts Wnt/β-catenin signaling. (A) Live imaging of embryos showed that in GFP and Nvβ-catenin-mCherry co-expressing embryos β-catenin-mCherry was enriched in the nuclei at one side of the embryo. However, co-overexpression of NvAxin-GFP (B) or SpAxin-GFP (C) with Nvβ-catenin-mCherry eliminated the nuclearization of β-catenin-mCherry. (D-F) Compared to GFP overexpressing control embryos which had a well-formed gut epithelium (D), NvAxin-GFP overexpressing embryos (E) and sea urchin Axin overexpressing embryos (SpAxin-GFP) (F) initiated primary invagination but failed to maintain a normal gut epithelium at the late gastrula stage. Green: phalloidin; Blue: DAPI. (G) Gene syn-expression domains as defined in Röttinger et al. (2012). Figure on the left shows a lateral view and the figure on the right shows the oral view. (H, I) qPCR analyses of NvAxin or SpAxin overexpressing embryos at 24 hpf (hours post fertilization) showed that all endomesoderm gene markers (orange, and red bars) and pharynx markers (blue bars) were downregulated, except for lhx6. The values represent the fold change relative to the GFP control (black bars). The qPCR experiments were replicated with three separate batches of embryos, with three technical replicates in each experiment. GAPDH was used as the internal control. Statistical significance was determined using the Unpaired t test. ****, p<0.0001; ***, p<0.001; **, p<0.01; *, p<0.05; n=3.

Earlier studies had demonstrated that overexpression of sea urchin Axin, NvDvl-DIX (a dominant-negative form of Dvl for cWnt signaling), a repressive form of β-catenin and dominant-negative NvTcf produced a similar phenotype in *Nematostella* embryos where the overexpression of the proteins blocked cWnt signaling, but primary archenteron invagination still occurred (Lee et al. 2007; Kumburegama et al. 2011; Röttinger et al. 2012). These studies noted that while the embryos in which cWnt was downregulated initially formed an endodermal epithelium, this epithelium appeared to disintegrate by the late gastrula stage (Kumburegama et al. 2011; Röttinger et al. 2012). Consistent with these previous observations, overexpression of NvAxin in embryos by mRNA injection into zygotes produced embryos that underwent primary archenteron invagination, and these embryos failed to maintain a normal gut epithelium compared to embryos expressing GFP (Fig. 2D-F).

We next investigated if manipulating Axin levels in *Nematostella* embryos had an effect on endogenous gene expression consistent with a function for the protein in the cWnt DC. We reasoned that overexpressing NvAxin would lead to a downregulation of cWnt-dependent gene expression while knockdown or knockout of NvAxin would lead to an increased expression of these genes. Consistent with these predictions, assaying NvAxin- or SpAxin-overexpressing *Nematostella* embryos for gene expression in syn-expression domains (Röttinger et al. 2012) (see Fig. 2G) using qPCR showed a downregulation of genes expressed in the central domain and central ring (see Fig. 2G) and an upregulation of genes expressed in the apical domain compared to control embryos (Fig 2H,I). Conversely, knockdown of NvAxin using antisense morpholinos led to an upregulation of genes expressed in the central domain, central ring, and external ring while genes expressed in the apical domain were downregulated (Supplementary Fig. 3A). To compare the changes in gene expression following knockdown of NvAxin with the changes in gene expression caused by knockdown of NvAPC-like, we injected antisense morpholinos against NvAPC-like into zygotes and used qPCR to analyze gene expression in these embryos. Consistent with the effect we noted in NvAxin knockdown embryos, we saw an upregulation of genes expressed in the central domain and the central ring in NvAPC-like MO injected embryos (Supplementary Fig. 3B).

To use an additional strategy to assess the effect of Axin downregulation on cWnt signaling in *Nematostella* embryos we carried out a CRISPR/Cas9-mediated NvAxin knockout experiment. To compare the molecular phenotypes between *NvAPC-like* and *NvAxin* knockout embryos we also designed an *NvAPC-like* single guide RNA (gRNA) to introduce a mutation into *NvAPC-like* in *Nematostella* embryos. The *NvAxin* and *NvAPC* gRNAs and Cas9 enzyme were injected into zygotes and animals were observed when control Cas9-only injected animals were at the four-tentacle polyp stage. We noted that *Nematostella* polyps developing from zygotes injected with *Axin* gRNA/Cas9 and *APC-like* gRNA/Cas9 had supernumerary tentacles (Kraus et al. 2016) (Fig. 3Aa, Ab, Ac). Animals injected with *Axin* plus *APC-like* gRNA with Cas9 typically showed a more severe phenotype with additional ectopic tentacles present throughout the animal (Fig. 3Ad). The NvAxin gRNA-mediated genome editing was validated by sequencing around the NvAxin gRNA excise sites in four single NvAxin gRNA-injected embryos and one Cas9-injected embryo (Supplementary Fig. 4) and by using WMISH to detect *NvAxin* expression in CRISPR/Cas9-mediated knockout embryos compared with Cas9-injected control embryos at 24 hpf (Fig. 3B). WMISH showed that 81.9% of the control embryos showed strong *NvAxin* expression at the animal pole; 13.7% showed detectable but weaker asymmetrical *NvAxin* expression, and 4.4% showed no detectable signal (n=204) (Fig. 3Ba). By contrast, 72.5% of NvAxin gRNA/Cas9-injected embryos showed no detectable *Axin* expression, 24.4% showed weak *Axin* expression, and only 3.1% showed a signal like that seen in the control (n=131) (Fig. 3Bb). The embryos with weak *NvAxin* signals may be due to incomplete excision of NvAxin or the generation of mosaic embryos where the gene was only excised in some nuclei. The percentage of successful excisions using CRISPR/Cas9 noted in this study are consistent with other studies that have used CRISPR/Cas9 to carry out gene editing in *Nematostella* embryos (Kraus et al. 2016; Servetnick et al. 2017).

**Figure 3.**
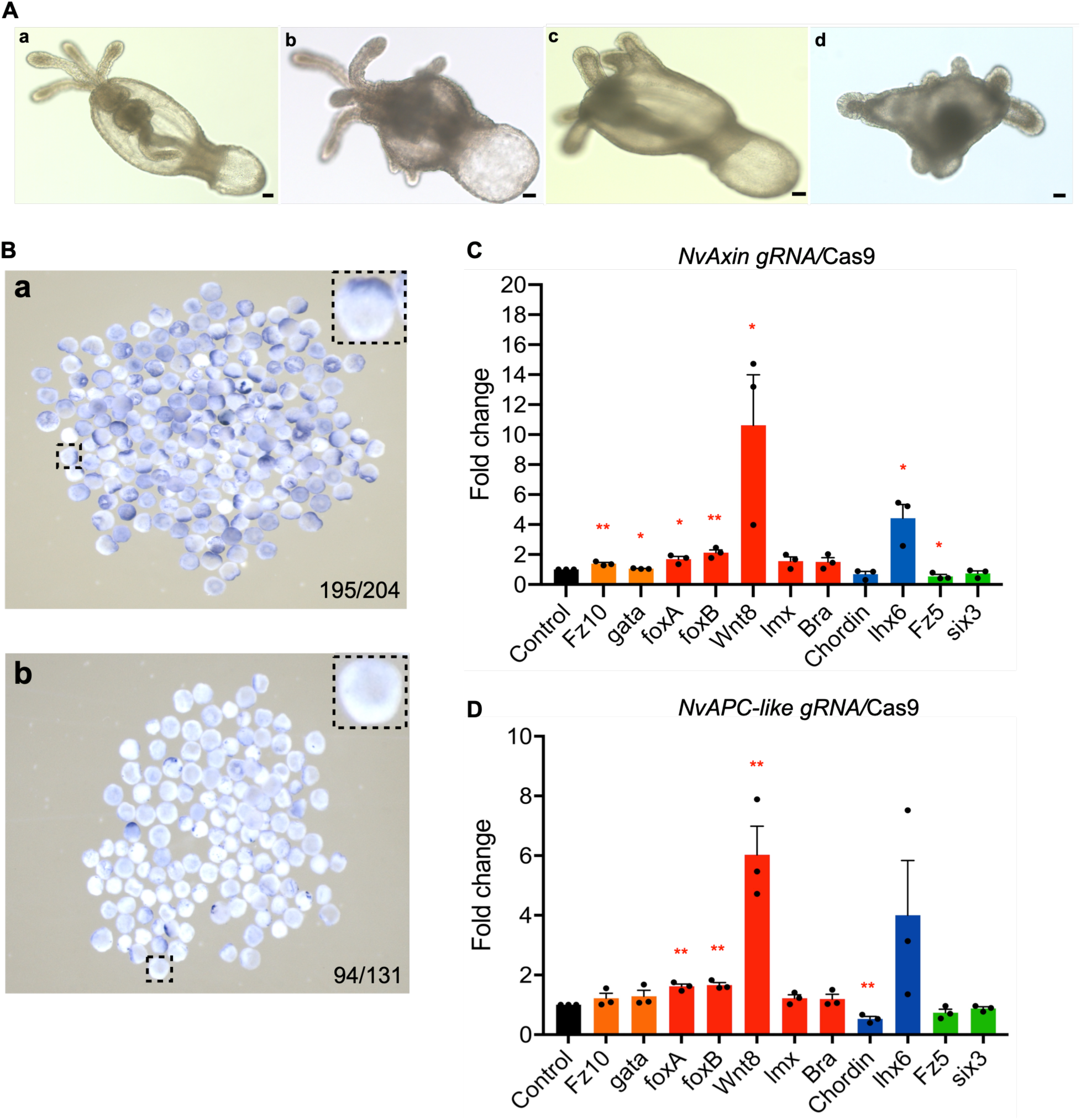
CRISPR/Cas9 mediated knockout of *Axin* and *APC-like* induces ectopic tentacles and upregulates endomesoderm gene expression in *Nematostella* embryos. (A) *Nematostella* 4-tentacle polyp, 7 days post-fertilization (7dpf) injected with Cas9 at the zygote stage (a). *NvAxin* guide RNA/Cas9-injected polyps produced ectopic tentacles (b), which is similar to what is seen in *NvAPC-like* gRNA/Cas9 injected polyps (c). Co-injection of *Axin* and *APC-like* gRNA/Cas9 produced a more severe phenotype (d) than *Axin* or *APC-like* gRNA/Cas9 single injections. Scale bar = 20µm. For each injection experiment, 100-200 embryos were injected with either gRNA/Cas9 or Cas9 alone. Among these, 70-75% of the gRNA/Cas9-injected embryos and more than 95% Cas9 only injected embryos exhibited the same morphology as shown in the representative figures. (B) While Cas9-injected embryos have normal oral pole *Axin* localization (a, blue staining), in situ hybridization showed that *NvAxin*/Cas9 injected embryos displayed reduced *Axin* expression (b). WMISH showed that 81.9% of the control embryos showed strong *NvAxin* expression at the animal pole; 13.7% showed detectable but weaker asymmetrical *NvAxin* expression, and 4.4% showed no detectable signal (n=204) (Fig. 2Ba). By contrast, 72.5% of *NvAxin* gRNA/Cas9-injected embryos showed no detectable *Axin* expression, 24.4% showed weak *Axin* expression, and only 3.1% showed a signal like that seen in the control (n=131) (Fig. 2Bb). The embryos with weak *NvAxin* signals may be due to incomplete excision of *NvAxin* or the generation of mosaic embryos where the gene was only excised in some nuclei. (C and D) qPCR shows the effects of *Axin* and *APC-like* knock-out by CRISPR/Cas9 on the expression of endomesoderm and pharynx gene markers. *Fz10*, *gata*, *foxA*, *foxB*, *Wnt8*, *lmx*, *Brachyury (Bra)* (orange and red bars) are endo mesodermal markers, *Chordin*, *lhx6* (blue bars) are markers for the pharynx; *Fz5* and *six3* (green bars) are ectoderm gene makers. qPCR experiments were replicated with three separate batches of embryos with three technical replicates in each experiment. GAPDH was used as the internal control. Graphs shown fold change in mRNA levels normalized to the Cas9 only control (black bars). Statistical significance was determined using the Unpaired t test. **, p<0.01; *, p<0.05; n=3.

To evaluate the effect of CRISPR/Cas9-mediated *Axin* and *APC-like* knockout on cWnt-dependent gene expression during early embryonic development, we used qPCR analysis and the results revealed that the genes expressed in the central domain and central ring were all upregulated indicating that knockout of *NvAxin* and *NvAPC-like* led to increased cWnt signaling consistent with a function for NvAxin and NvAPC-like in the cWnt DC (Fig. 3C,D). The qPCR analysis also revealed that the apical domain genes *Fz5* and *Six3* were downregulated, consistent with previous studies showing that global upregulation of cWnt signaling leads to the downregulation of genes expressed in the apical domain (Röttinger et al. 2012) (Fig. 3C,D). Notably, *NvWnt8* and *Nvlhx6* (a gene expressed in the external ring) exhibited a stronger upregulation in both *NvAxin* gRNA and *NvAPC-like* gRNA-injected embryos than other genes expressed at the animal pole, suggesting that these genes are particularly sensitive to levels of cWnt signaling. Hence, disruption of either one of the two core scaffold proteins in the *Nematostella* DC likely resulted in the strong activation of *Wnt8* and *lhx6* expression. Alternatively, *NvWnt8* and *Nvlhx6* might be repressed by cWnt outside their normal domain of expression and are ectopically expressed when cWnt is downregulated. In sum, these results provided strong evidence that Axin plays a crucial role in negatively regulating cWnt signaling in *Nematostella* embryos and indicated that NvAxin has a role in the cWnt DC even though it lacks a distinct β-catenin binding motif. Additionally, these experiments showed that NvAPC-like regulates cWnt signaling even though it lacks most of the domains shown be required for APC function in the DC in bilaterians. Our work also corroborated the work of Kraus et al. (2016) who previously showed that NvAPC-like regulates cWnt signaling in *Nematostella*.

### Nematostella Axin overexpression disrupts cWnt signaling in sea urchin embryos

We next sought to determine if *Nematostella* Axin could regulate cWnt signaling in a bilaterian. Extensive work done in sea urchin embryos has shown that the cWnt pathway plays a critical role in specifying and patterning the anterior-posterior (AP) axis, and blocking β-catenin nuclearization in early embryos produces a unique anteriorized phenotype hence making these embryos useful for assaying cWnt signaling (Wikramanayake et al. 1998; Logan et al. 1999; Sun et al. 2021). When we expressed sea urchin β-catenin-mCherry (Spβ-catenin-mCherry) in sea urchin embryos by mRNA injection into zygotes, live imaging showed typical nuclearization of Spβ-catenin-mCherry at the vegetal pole (Wikramanayake et al. 1998; Logan et al. 1999; Sun et al. 2021) (Fig. 4Aa). However, when Spβ-catenin-mCherry was co-expressed with NvAxin or SpAxin in sea urchin embryos there was complete elimination of Spβ-catenin-mCherry nuclearization (Fig. 4Ab, Ac). Moreover, consistent with previous observations in sea urchin embryos where blocking Spβ-catenin nuclearization by SpAxin overexpression led to severe anteriorization of sea urchin embryos (Sun et al. 2021) (Fig. 4Ba,Bb), overexpression of NvAxin in sea urchin embryos also led to a phenotype indistinguishable from the phenotype generated from SpAxin overexpression (Fig. 4Bc). Importantly, the anteriorized embryos were not able to recover even when control animals were at the pluteus larval stage (Fig. 4Ba’-c’) (Wikramanayake et al. 1998; Sun et al. 2021). Consistent with the anteriorized phenotype, analysis of gene expression in sea urchin embryos overexpressing NvAxin and SpAxin showed that all seven endomesoderm gene markers tested were significantly downregulated, while two neuroectoderm gene markers were upregulated (Fig. 4C, D). These experiments demonstrated that NvAxin is a strong regulator of cWnt signaling in a bilaterian, even though it lacks the conserved βcatBM.

**Figure 4.**
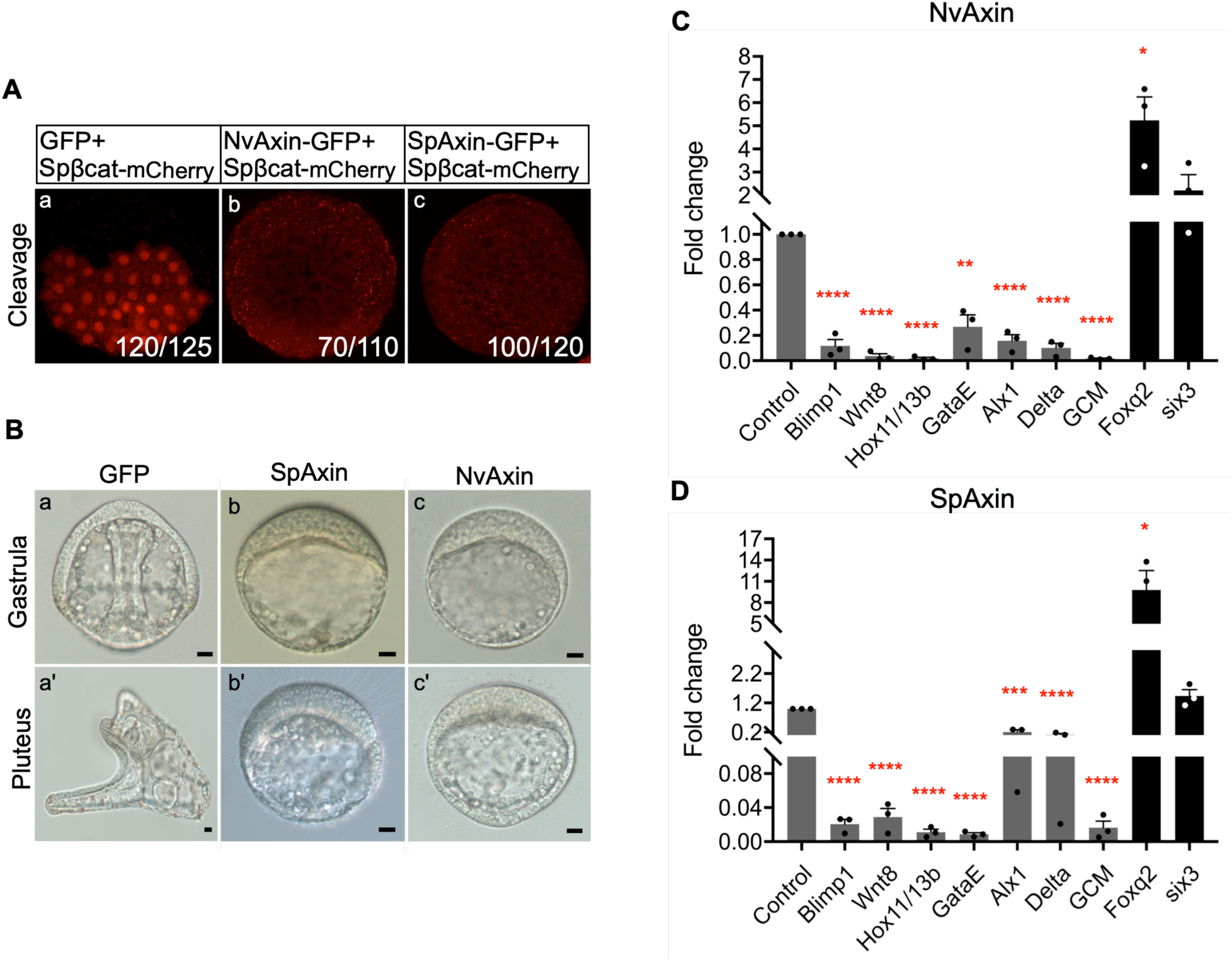
Overexpression of *Nematostella* Axin in sea urchin embryos blocks nuclearization of β-catenin and cWnt signaling. (A) In the control where GFP and Spβ-catenin-mCherry were co-expressed, β-catenin was enriched in the nuclei at the vegetal pole (a). Co-overexpression of NvAxin (b) or SpAxin (c) with Spβ-catenin-mCherry downregulated the nuclearization of β-catenin. (B) Control embryos injected with *GFP* mRNA developed normally (a, a’), while embryos developing from *SpAxin* or *NvAxin* mRNA-injected zygotes became severely anteriorized (b, b’ and c, c’). Scale bar = 10µm. (C and D) qPCR analyses showed that NvAxin or SpAxin overexpressing embryos downregulated endomesoderm genes and upregulated neuroectoderm genes. The bar graph shows the fold change of each gene between *Axin* mRNA-injected and *GFP* mRNA-injected control embryos. *Blimp1*, *Wnt8*, *Hox11/13b*, *GataE*, *Alx1*, *Delta* and *GCM* are endomesoderm gene markers; *Foxq2* and *Six3* are neuroectoderm gene markers. qPCR experiments were replicated with three separate batches of embryos with three technical replicates in each experiment. GAPDH was used as the internal control. Statistical significance was determined using the Unpaired t test. ****, p<0.0001; ***, p<0.001; **, p<0.01; *, p<0.05; n=3. For each injection experiment, 200-300 embryos were injected with each construct and more than 95% of embryos had the same morphology as shown in the representative figures.

### The NvAxin GSK-3β binding domain is required for its function in the cWnt pathway

In bilaterians, the βcatBM on Axin is critical for the protein to interact with and target β-catenin for degradation during activation of cWnt signaling (Ikeda et al. 1998; Xing et al. 2003; Stamos and Weis 2013). Bioinformatic analysis using SMART indicated that NvAxin lacks the βcatBM but it has the APC- (RGS), GSK- (GID), and Dvl- (DAX) binding domains, which are also found in the Axin proteins of other non-bilaterians (Supplementary Table 1). Since our studies showed that NvAxin could regulate β-catenin nuclearization in the sea urchin embryo, we used a rescue assay that we had previously developed in sea urchins (Sun et al. 2021) to identify the domains on NvAxin required for β-catenin regulation in the cWnt pathway. This rescue assay takes advantage of the unique “posteriorized” phenotype produced in sea urchin embryos when levels of Axin in the embryo are reduced using an anti-sea urchin Axin morpholino, which can be rescued by overexpressing the wildtype Axin protein (Sun et al. 2021) (Fig. 5A-C). We showed that while injection of a control MO has no effect on embryo development, injections of an anti-sea urchin (*Lytechinus variegatus*, Lv) Axin MO produces a posteriorized phenotype, and additionally, while *GFP* RNA control injections have no effect on development, *NvAxin* RNA injection anteriorizes sea urchin embryos (Fig. 5Ba-d’). To determine if NvAxin could rescue sea urchin embryos posteriorized by injection of the LvAxin MO, we co-injected the LvAxin MO with full-length *NvAxin* mRNA. Examination of these embryos showed that while the embryos co-injected with the LvAxin MO and *GFP* mRNA were posteriorized as expected (Fig. 5Ca,a’), the embryos co-injected with the LvAxin MO and *NvAxin* mRNA were completely rescued (Fig. 5Cb,b’). Similarly, co-injection of the LvAxin morpholino with *NvAxinΔRGS* mRNA or *NvAxinΔDAX* mRNA efficiently rescued the posteriorized phenotype induced by the LvAxin morpholino in sea urchins (Fig. 5Cc-d’). However, co-injection of the LvAxin MO with *NvAxinΔGID* mRNA failed to rescue the phenotype (Fig. 5Ce,e’) indicating the requirement of the GSK-3β binding domain (GID) for NvAxin to function in the cWnt DC. This result is consistent with our observations in sea urchin embryos, where the SpAxinΔGID construct was also unable to rescue Axin knockdown embryos (Sun et al. 2021). We conclude that the GID domain is likely the most crucial domain of NvAxin for regulating β-catenin stability in the cWnt DC.

**Figure 5.**
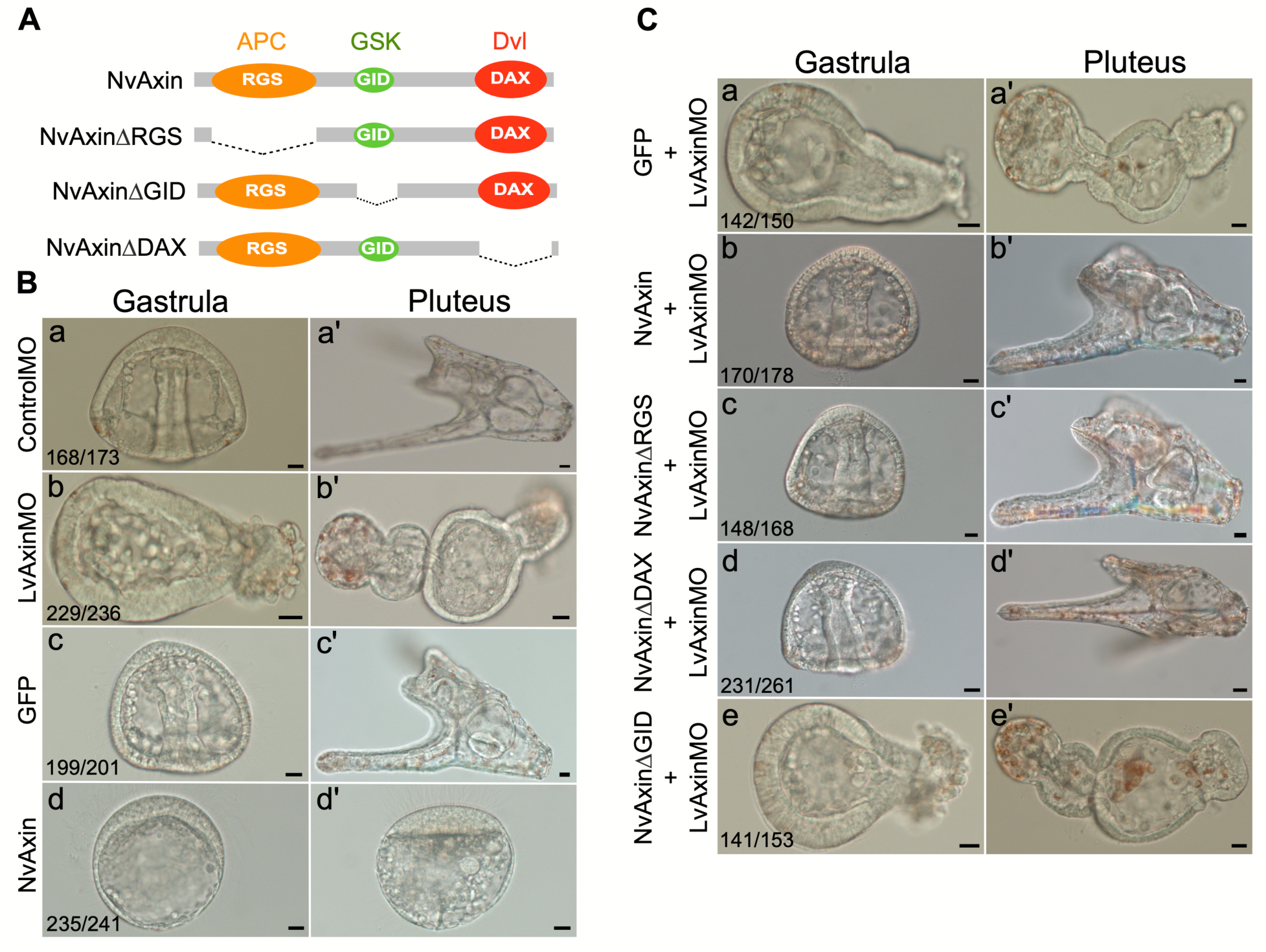
Sea urchin embryos posteriorized by Axin knock-down can be rescued by full-length NvAxin, NvAxinΔRGS, and NvAxinΔDAX, but not NvAxinΔGID. (A) Schematic representations of full-length Axin and single domain-deleted Axin constructs. (B) Gastrula and pluteus larva developing from Control MO, LvAxin MO, *GFP* mRNA or *NvAxin* mRNA injected zygotes from the sea urchin *Lytechinus variegatus* (a-d’). Control MO-injected embryos develop normally (a,a’), while LvAxin MO-injected embryos become posteriorized (b, b’). *GFP* mRNA-injected zygotes produce normal embryos (c, c’), while *NvAxin* mRNA-injected zygotes develop into anteriorized embryos (d, d’). (C) The posteriorized phenotype induced in LvAxin MO-injected embryos is not rescued by control *GFP* mRNA co-injection (a, a’) but is rescued by the overexpression of NvAxin (b, b’), NvAxinΔRGS (c, c’), and NvAxinΔDAX (d, d’). Injection of *NvAxinΔGID* mRNA fails to rescue the posteriorization induced by Axin knockdown (e,e’). Scale bar = 10 µm. The numbers in the panels indicate the number of embryos examined and the number that showed the phenotype.

Previous studies in bilaterians have shown that overexpression of the Axin GID peptide alone is sufficient to activate cWnt signaling (Hedgepeth et al. 1999; Sun et al. 2021). The underlying mechanism for this is still unclear but it has been suggested that the GID domain can bind to endogenous GSK-3β and prevent it from interacting with Axin (Hedgepeth et al. 1999; Chou et al. 2006; Sun et al. 2021). Our NvAxin rescue experiments in sea urchins using Axin domain-deletion constructs clearly showed that the GID domain is required for NvAxin’s activity in the cWnt destruction complex (Fig. 5Ce,e’). GSK-3β binding to Axin is a critical step in assembling the DC in bilaterians. To begin to assess if the NvAxin-GID (NvGID) domain was involved in regulating GSK-3β in *Nematostella* we first overexpressed an NvGID-GFP construct in embryos by microinjection of mRNA into zygotes. When control animals were at the polyp stage, the NvGID-GFP overexpressing embryos had a morphology resembling *Nematostella* embryos overexpressing an activated form of β-catenin (Röttinger et al. 2012) (Fig. 6A). Analysis of germ layer-specific markers at the blastula stage using qPCR showed that endomesoderm gene markers and pharyngeal markers were highly upregulated while the ectoderm gene markers were downregulated (Fig. 6B). These results are consistent with the notion that as seen in bilaterians, Axin-GID overexpression can activate cWnt signaling and provides additional evidence that NvAxin scaffolds a critical DC component similar to bilaterian Axin.

**Figure 6.**
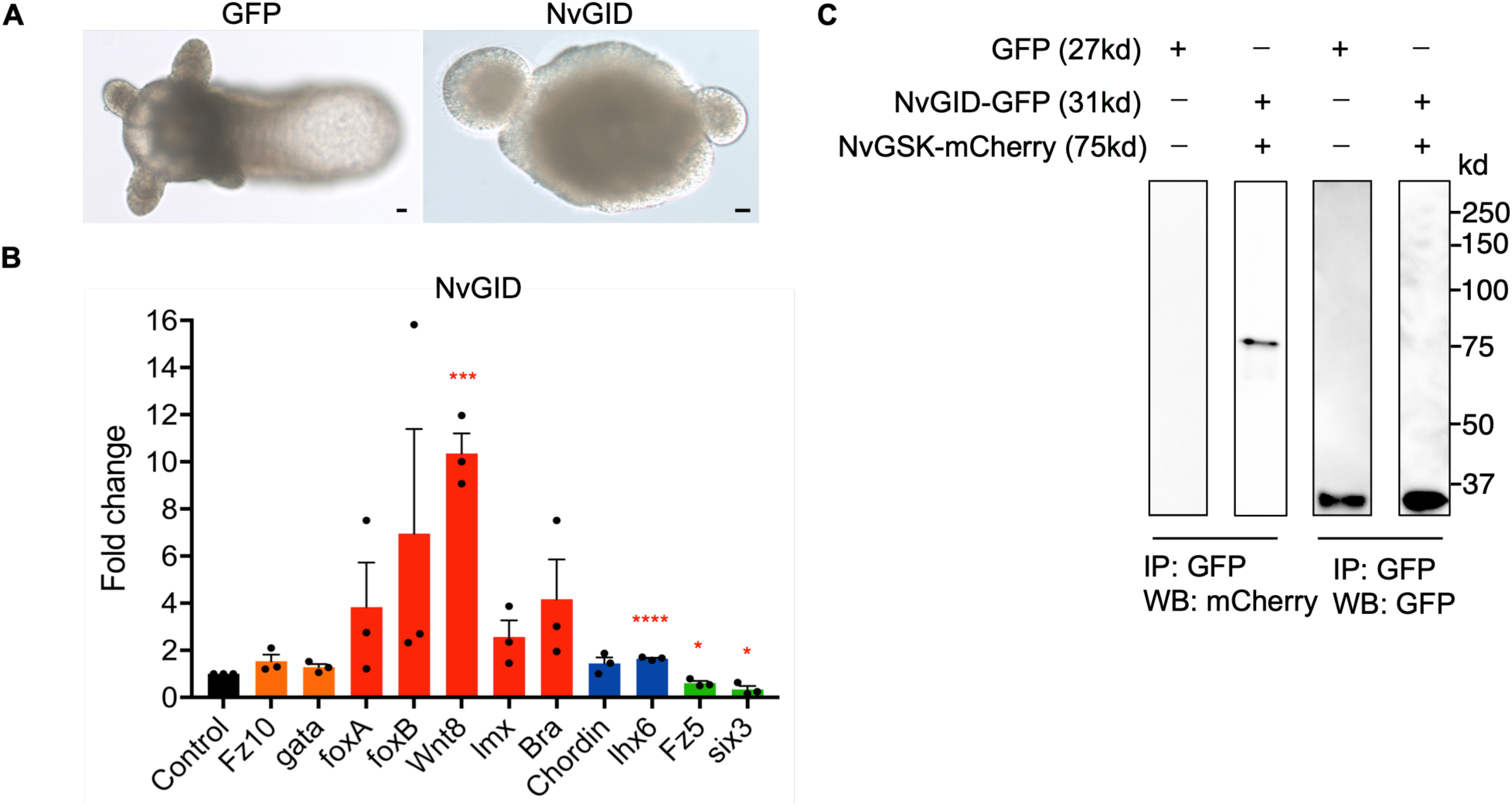
Overexpression of the NvAxin GID domain upregulated endomesodermal and pharyngeal gene expression in *Nematostella* embryos. (A) Comparison of animals developing from *Nematostella* zygotes injected with *GFP* mRNA (control) or *NvAxin-GID (NvGID)* mRNA at 7 days post fertilization (7dpf). At 7dpf control animals were at the polyp stage while *NvGID* RNA-injected embryos developed abnormally. Scale bar =10 µm. (B) qPCR analyses of NvGID overexpressing embryos at 24 hpf (hours post fertilization) showed that all endomesoderm gene markers (orange, red and blue bars) were upregulated. Some genes were unregulated at modest levels while others showed significant increases. The two apical domain genes were significantly downregulated. The qPCR experiments were replicated with three separate batches of embryos, with three technical replicates in each experiment. GAPDH was used as the internal control. Graphs shown fold change in mRNA levels normalized to the GFP control (black bars). Statistical significance was determined using the Unpaired t test. ****, p<0.0001; ***, p<0.001; *, p<0.05; n=3. (C) Western blot analysis of anti-GFP antibody-mediated immunoprecipitation of *in vitro* synthesized NvGID-GFP and NvGSK-3β-mCherry fused proteins.

The residues within the GID domain mediating interaction between Axin and GSK-3β are highly conserved in bilaterians and all four non-bilaterian taxa (Kremer et al. 2010) (Supplementary Fig. 5). The conservation of the NvGID suggested that the phenotype elicited by overexpression of the peptide in *Nematostella* was due to it directly binding to NvGSK-3β and preventing NvGSK-3β from binding to NvAxin. To validate this predicted interaction, we mixed NvGID-GFP and NvGSK-3β-mCherry proteins made using *in vitro* transcription and translation (TnT) and immunoprecipitated (IP) NvGID-GFP using anti-GFP antibodies. Western blot analysis of the IP samples using anti-mCherry antibodies showed a clear interaction between NvGID-GFP and NvGSK-3β-mCherry *in vitro* (Fig. 6C). These results demonstrated that NvGID directly interacts with NvGSK-3β and further supports the notion that NvAxin has an activity consistent with it being part of a DC in *Nematostella*.

It is important to note that the morphological and molecular phenotypes generated from upregulating cWnt signaling by overexpression of NvGID, and NvAxin and NvAPC-like knock down/knock out using morpholinos and CRISPR/Cas-9 show some significant differences (compare Fig. 3 with Fig. 6). When NvAxin-GID was overexpressed in *Nematostella* embryos, the embryos did not form a polyp stage like controls, but rather had what appeared to be a hollow and expanded swollen appearance (Fig. 6A). Gene expression analysis showed that these embryos had elevated expression of genes expressed in the central domain, central ring and external ring and reduction in expression of genes expressed in the apical domain. These embryos resemble the phenotypes generated when ectopically activating cWnt signaling by overexpressing a constitutively activate form of β-catenin (Röttinger et al. 2012). Both these methods are likely to activate early cWnt signaling thus impacting early cell fate specification in *Nematostella* embryos. It is likely that there is Axin and APC-like mRNA and protein in the egg and early embryo and hence, when using morpholinos and CRISPR/Cas-9 to knock down or knock out Axin and APC-like, these methods do not impact the maternally loaded molecules. We suggest that the Axin and APC-like knock-down/knock-out phenotypes are due to elevated cWnt signaling affecting oral-aboral axis patterning, a relatively late event, and not early cell fate specification.

### In vitro and in vivo protein-protein interaction assays indicate that the Nematostella Destruction Complex proteins have binding activities previously undefined in bilaterians

Previous studies in bilaterian taxa such as vertebrates and dipterans have shown interactions between Axin-APC, Axin-β-catenin, APC-β-catenin, APC-GSK-3β and Axin-GSK-3β *in vitro* and *in vivo* and in general, the protein binding interfaces for these interactions have been well defined (Ikeda et al. 1998; Kishida et al. 1998; Dajani et al. 2003; Ha et al. 2004; Peterson-Nedry et al. 2008; Kremer et al. 2010; Roberts et al. 2011; Pronobis et al. 2015; Ranes et al. 2021). These studies have also demonstrated that while there is a single defined β-catenin binding domain on bilaterian Axin, there are multiple β-catenin binding domains on APC that have distinct sequence motifs (Roberts et al. 2011; Pronobis et al. 2015; van Kappel and Maurice 2017; Pronobis et al. 2017). The experimental analyses that we have done in this study showed that both NvAxin and NvAPC-like regulate cWnt in *Nematostella*. Hence, the lack of critical domains required for cWnt DC activity in NvAxin and NvAPC-like prompted us to investigate if these proteins could still form a functional DC perhaps through protein binding domains previously uncharacterized in bilaterians. To explore this possibility, we synthesized GFP-tagged NvAxin and SpAxin, and mCherry-tagged Nvβ-catenin, Spβ-catenin, NvGSK-3β and SpGSK-3β using *in vitro* TnT and then mixed the various protein pairs and used an anti-GFP antibody to IP the GFP tagged Axin proteins. We then probed Western blots of the immunoprecipitated proteins with anti-mCherry antibodies to evaluate any interaction between the protein pairs. As expected, this assay revealed a robust interaction between sea urchin Axin-GFP and sea urchin β-catenin-mCherry (Fig. 7A, Lane 5). We also noted a weaker but persistent interaction between NvAxin and Nvβ-catenin and NvAxin and Spβ-catenin proteins in vitro (Fig. 7A, Lane 1,2). This was unexpected since NvAxin does not have the well-defined βcatBM seen in bilaterian Axin (Fig. 1C).

**Figure 7.**
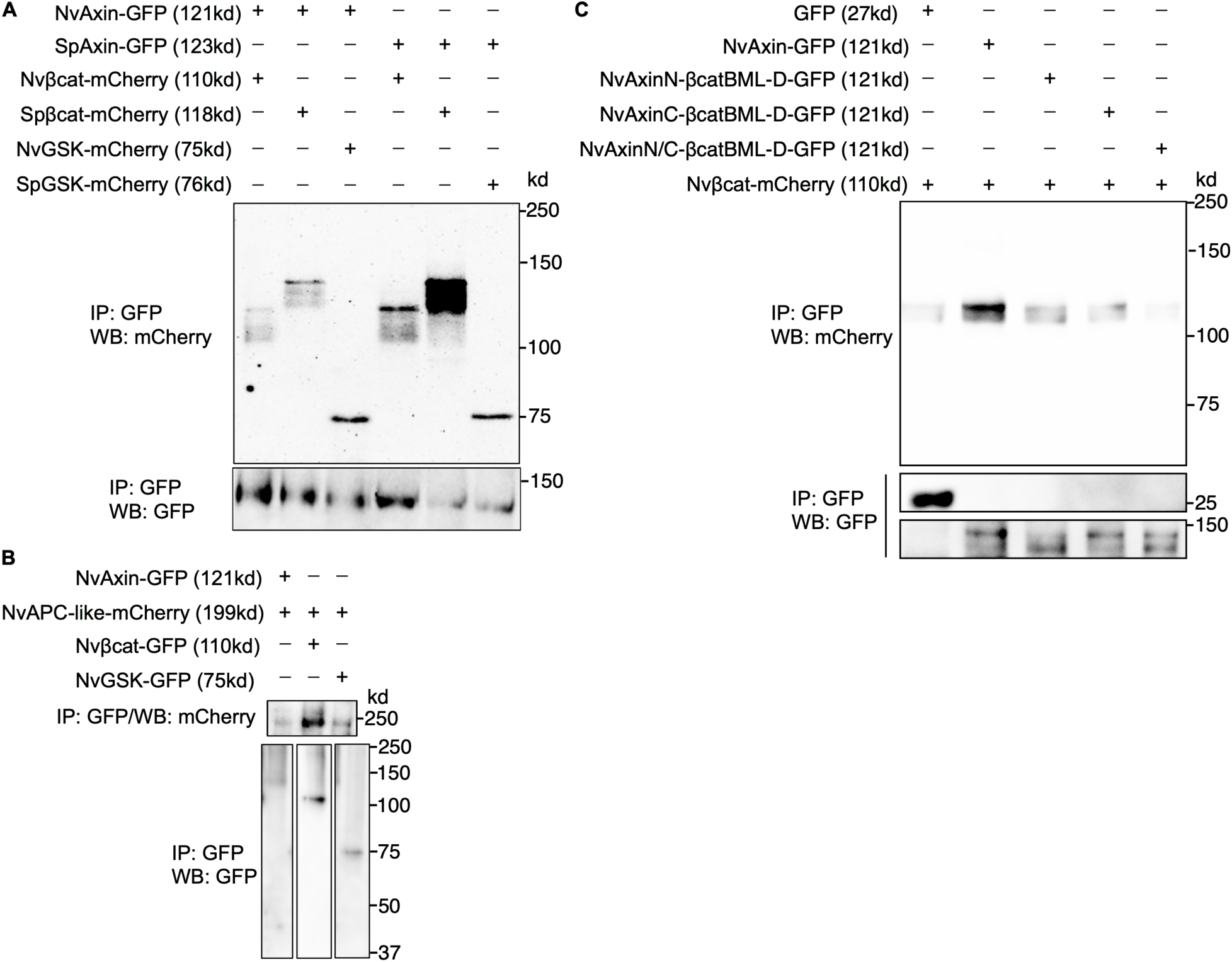
*In vitro* analysis of protein-protein interactions indicate direct interactions between Nvβ-catenin and components of the *Nematostella* β-catenin destruction complex. (A) *In vitro*-synthesized GFP- or mCherry-fused proteins were tested for potential interactions by mixing a GFP-fused partner with an mCherry-fused partner and performing immunoprecipitation (IP) with an anti-GFP antibody. The labels at the top of the panel show the interacting proteins in each experiment. The top part of the Western blot analysis shows the protein-protein interactions between protein pairs. The bottom part of the figure shows the Western blot of GFP-tagged proteins. Sea urchin (*Strongylocentrotus purpuratus*) β-catenin (Spβ-catenin) shows a strong interaction with sea urchin Axin (SpAxin) (Lane 5), while Nvβ-catenin shows a consistent but considerably weaker interaction with NvAxin (Lane 1). NvAxin also shows a relatively weaker interaction with Spβ-catenin (compare lanes 2 and 5). Unexpectedly, SpAxin shows a weaker interaction with Nvβ-catenin than with SpAxin (compare lane 4 and lane 5). Sp and Nv Axin bind the SpGSK-3β and NvGSK-3β respectively (Lanes 3 and 6). (B) NvAPC-like and Nvβ-catenin interact strongly in vitro (Lane 2) while NvAPC-like shows weaker but consistent interactions with NvAxin and NvGSK-3β (Lanes 1 and 3). (C) NvAxin-GFP and Nvβ-catenin-mCherry interact in vitro (Lane 2) as previously shown (Figure 7A, Lane 1), changing the conserved leucine on the Axin-RGS located βcatBM-like sequence to aspartic acid reduces the in vitro interaction between NvAxin-GFP and Nvβ-catenin-mCherry (Lane 3) as does the same mutation to the βcatBM-like sequence towards the C-terminus of NvAxin (Lane 4). Mutating the leucine to aspartic acid in both NvAxin βcatBM-like sequences completely abolishes the interaction of NvAxin-GFP with Nvβ-catenin-mCherry (Lane 5).

We next asked if sea urchin Axin (which has the conserved βcatBM) could interact with Nvβ-catenin. To our surprise, despite sea urchin Axin containing a conserved βcatBM and a demonstrated ability to strongly interact with sea urchin β-catenin *in vitro*, it only displayed a weaker interaction with Nvβ-catenin in this assay (Compare Fig. 7A, Lane 4 and Lane 5). These results suggested that the Axin interacting domains on Nvβ-catenin (the armadillo repeats) had diverged from the corresponding domains on bilaterian β-catenin, and hence it was unable to interact strongly with a bilaterian Axin. *Nematostella* β-catenin has all 12 ARM domains seen in bilaterians and each of its ARM repeats has a high level of homology to corresponding ARM domains in bilaterian β-catenin indicating a strong conservation over 700 million years of evolution (Schneider et al. 2003). Hence, any changes in Nvβ-catenin protein that affects how it interacts with a bilaterian Axin βcatBM would likely be more subtle amino acid changes that affects binding affinity between the proteins. In sum, these experiments showed that NvAxin can directly interact with *Nematostella* and sea urchin β-catenin, but based on the *in vitro* assay, these interactions appeared considerably weaker than the interaction between sea urchin Axin and sea urchin β-catenin.

We then investigated if NvAxin could directly interact with NvGSK-3β using the *in vitro* pull-down assay. As predicted by the presence of a conserved GID on NvAxin and our previous observation that the NvGID could interact with NvGSK-3β, Western blotting of the IP products showed a clear interaction between NvAxin-GFP and NvGSK-3β-mCherry (Fig. 7A, Lane 3). Likewise, our results showed that sea urchin Axin-GFP interacted strongly with sea urchin GSK-3β-mCherry (Fig. 7A, Lane 6). It is notable that the Axin GID domain is highly conserved, including in ctenophores (Supplementary Fig. 5), indicating that the Axin-GSK-3β interaction evolved over 700 mya. In bilaterians, Axin and APC proteins heterodimerize to form the scaffold for the β-catenin DC. Bilaterian APC/APC-like proteins have a complex domain structure that allows them to interact with several proteins including Axin and β-catenin. NvAPC-like is smaller in size than the vertebrate or dipteran proteins, and it lacks many of the domains shown to bind to β-catenin and Axin in bilaterians (Fig. 1D; Roberts et al. 2011; Pronobis et al. 2015; Pronobis et al. 2017; Ranes et al. 2021). When we analyzed NvAPC-like domain structure using the webtool SMART (Simple Modular Architecture Research Tool), we noted that it had seven conserved armadillo repeats and two poorly characterized repeat regions at the N-terminus. NvAPC-like also lacked the SAMP domains that are critical for bilaterian Axin-APC interaction (Pronobis et al. 2015; Pronobis et al. 2017) (Fig. 1D). To determine if NvAPC-like (tagged with mCherry) could directly interact with NvAxin, Nvβ-catenin and NvGSK (tagged with GFP), we carried out the *in vitro* protein-protein interaction assay previously described. We noted that NvAPC-like-mCherry robustly interacted with Nvβ-catenin-GFP in this assay (Fig. 7B). We were also able to detect interactions between NvAPC-like-mCherry and NvGSK-3β-GFP and NvAPC-like-mCherry and NvAxin-GFP *in vitro* (Fig. 7B).

The *in vitro* assay that we used in the experiments described above provides insight into potential direct interactions between components of the cWnt DC in *Nematostella* embryos. However, in those cases where we saw weak interactions *in vitro*, it is possible that in the *in vivo* environment, post-translational modifications or bridging by other proteins in the embryo lead to higher affinity interactions (Ha et al. 2004; Kimelman and Xu 2006; Pronobis et al. 2015; Ranes et al. 2021). To validate the results that we saw in the *in vitro* experiments with an *in vivo* and *ex vivo* approach, we adopted the split Click Beetle Green (CBG) luciferase system (Villalobos et al. 2010) (Supplementary Fig. 6). Axin and β-catenin proteins from *Nematostella* and the sea urchin *S. purpuratus* were tagged with split CBG luciferase (Villalobos et al. 2010). All possible pairs of split-luciferase-tagged Axin and β-catenin proteins from the same or different species, were expressed in *Nematostella* embryos by mRNA injection into zygotes. The luminescence of split luciferase was measured in live embryos or in embryo lysates. Normalized kinetic curves from embryonic lysates and live embryos are shown in Supplemental Fig. 6. Consistent with the *in vitro* studies, SpAxin + Spβ-catenin had the highest signal in both embryonic lysates and live embryos, while NvAxin + Spβ-catenin had clear but relatively lower signal (Supplemental Fig. 6A, B). However, SpAxin did not appear to interact with Nvβ-catenin in either embryonic lysates or live embryos (11 % and 10 % of SpAxin + Nvβ-catenin signal, respectively). This result is inconsistent with the *in vitro* pull-down experiments where we noted a weak interaction between SpAxin and Nvβ-catenin (Fig. 7, Lane 4). The reason for this discrepancy is not known at this time. Interestingly, NvAxin + Nvβ-catenin yielded a relatively low signal (27% of NvAxin + Nvβ-catenin intensity) in live embryos but virtually none in lysates (13 %). These observations suggest that this interaction requires the presence of intact cellular and molecular complexes that are lost in lysates. However, overall, the interactions between NvAxin + Nvβ-catenin, NvAxin + Spβ-catenin, and SpAxin + Spβ-catenin were consistent between the *in vitro* and *in vivo* approaches and gives credence to the notion that the interaction between NvAxin and Nvβ-catenin is weaker than the interaction between SpAxin and Spβ-catenin. Moreover, the *in vivo* and *ex vivo* experiments confirmed that NvAxin and Nvβ-catenin, and SpAxin and Nvβ-catenin are only capable of relatively weak interactions most likely discounting a major *in vivo* role for bridging proteins or post-translational modifications to enhance these interactions.

### AlphaFold software predicts novel protein-binding interfaces between cWnt Destruction Complex components in Nematostella

The unexpected direct protein-protein interactions we noted in the *in vitro* assays prompted us to use AlphaFold3 to determine if the AI-based program would be able to predict the binding interfaces between the respective protein pairs. As expected, AlphaFold analysis predicted the interactions between SpAxin-Spβ-catenin and SpAxin-SpGSK-3β based on complementary domains seen in orthologs from other bilaterian proteins (Supplementary Fig. 7A, B). Moreover, as expected from the conserved domains, AlphaFold also predicted an interaction between NvAxin and NvGSK-3β (Supplementary Fig. 7C). The AlphaFold analysis also indicated possible interactions between NvAxin-Nvβ-catenin and NvAPC-like-Nvβ-catenin based on the iPTM and PTM values (Supplementary Fig. 7D, E). This was unexpected because NvAxin and NvAPC-like lack the β-catenin-interaction domains extensively characterized in bilaterians (Roberts et al. 2011; Pronobis et al. 2015; Pronobis et al. 2017; Ranes et al. 2021). In contrast, even though the *in vitro* assays showed interactions between NvAxin-NvAPC-like and NvAPC-like-NvGSK-3β, these interactions were not supported by AlphaFold analysis. (Supplementary Fig. 7F, G). This may be due to the NvAPC-like being a relatively large protein and in some such cases AlphaFold is not as efficient in predicting protein-protein interactions (Bryant et al. 2022; Shor and Schneidman-Duhovny 2024).

When we used AlphaFold3 and ChimeraX (Meng et al. 2023) to analyze predicted and visualized interactions sites between the NvAPC-like and Nvβ-catenin, we identified an α-helix interaction site within the ARM 708-753aa region (first ARM domain) of NvAPC-like. This region was predicted to interact with Nvβ-catenin with high iPTM and PTM values (Supplementary Fig. 7H). Notably, this ARM sequence is conserved across *H. sapiens*, *D. melanogaster*, *S. purpuratus* and *N. vectensis* (amino acid identity > 41%). We did not analyze the potential interactions of the other six NvAPC-like ARM domains with Nvβ-catenin. Finally, we analyzed NvAPC-like protein structure for a potential interaction with the NvAxin-RGS domain and AlphaFold strongly predicted an interaction between the NvAxin-RGS and Repeat 1 of NvAPC-like (Supplementary Fig. 7I) but not Repeat 2 (not shown).

The persistent, albeit weak, *in vitro* and *in vivo* interactions we noted between NvAxin and Nvβ-catenin (Fig. 7, Supplementary Fig. 6) and the prediction of an interaction between these proteins by AlphaFold3 analysis (Supplementary Fig. 7D) were unexpected. AlphaFold 3 and ChimeraX software predicted two possible regions of interaction between NvAxin and Nvβ-catenin, one located within the NvAxin-RGS domain, and the other closer to the C-terminus of NvAxin (Figure 7C; Supplementary Fig. 8A,B). Alignment of the predicted NvAxin-Nvβ-catenin interaction site on NvAxin with the highly conserved βcatBM in bilaterian Axins using Clustal Omega and Jalview showed some highly conserved amino acids between these sequences (Supplementary Fig. 8F). The amino acid identity between the sequence in NvAxin-RGS and the second sequence closer to the C-terminus, and the bilaterian βcatBM ranged from 20-26% and 11-28% respectively and we designated these sequences as “βcatBM-like”. Significantly, the leucines at positions 119 and 434 in the βcatBM-like sequences corresponds to a leucine residue known to be essential for the interaction between bilaterian Axin and β-catenin (Kremer et al. 2010) (Supplementary Fig. 8F). Additionally, Kremer et al. (2010) identified a histidine residue at position 504 in *Drosophila* Axin as being critical for β-catenin interaction with Axin. In *Nematostella* Axin, this corresponding residue is missing, possibly accounting for the weak interaction between NvAxin and Nvβ-catenin. We then examined Axin sequences from several other cnidarians to ask if similar duplicated βcatBM-like sequences were present in the Axin proteins. Interestingly, the predicted Axin-β-catenin binding sites are all located within the RGS domain (Supplementary Fig. 9A) and C-terminal (Supplementary Fig. 9B) of Axin and are highly conserved in each of these taxa. We also carried out AlphaFold and ChimeraX analysis of predicted interactions between Axin and β-catenin in *Tricoplax adhaerens* (a placozoan), *A. queenslandica* (a poriferan) and *Mnemiopsis leidyi* (a ctenophore). This analysis predicted a single putative interaction site between Axin and β-catenin in these species (Supplementary Fig. 8C,D,E,F). Alignment between the identified βcatBM-like sequence on the ctenophore MlAxin-RGS and the bilaterian βcatBM did not reveal conservation of the leucine residue; instead, an aspartic acid was identified (Supplementary Fig. 8F). However, the identified βcatBM-like sequences on AqAxin-RGS and TaAxin-RGS retained the conserved leucine associated with functional binding (Kremer et al. 2010) (Supplementary Fig. 8F). The amino acid identities between the βcatBM-like sequences in the RGS domain of TaAxin, AqAxin and MlAxin, and the bilaterian βcatBM ranged from 16-33%, 12-25% and 21-28% respectively. We used AlphaFold software to determine if the RGS domains of bilaterian Axins, including all the bilaterian species listed in Supplementary Table 1, contained any putative β-catenin interaction motifs, but we were not able to detect any such sequences (not shown).

Ctenophores are now widely believed to be the earliest-emerging metazoan group (Dunn et al. 2008; Schultz et al. 2023). Consistent with this notion, the βcatBM-like sequence on MlAxin identified using AlphaFold3 and ChimeraX lacked the conserved leucine residue that is present in Axins from the other three non-bilaterian taxa. This leucine along with a conserved histidine have been implicated as the two critical amino acids involved in Axin-β-catenin interaction in bilaterians. The absence of both the leucine and the histidine from the βcatBM-like sequence in ctenophore Axin indicated that it would not be able to directly interact with ctenophore β-catenin. Consistent with this idea MlAxin and Mlβ-catenin did not show any direct interactions when tested using the *in vitro* TnT assay (Supplementary Fig. 8G). We next addressed if the conserved leucine residues seen in the two βcatBM-like sequences in NvAxin were responsible for the “weak” binding of NvAxin and Nvβ-catenin. We used site-directed mutagenesis to change the conserved leucine in each of the βcatBM-like sequences in NvAxin to aspartic acid to resemble the ctenophore βcatBM-like sequence and tested the ability of these mutant NvAxin proteins to interact with Nvβ-catenin. Our analysis showed that mutating the conserved leucine on each of the two βcatBM-like sequences on NvAxin to aspartic acid reduced interaction with Nvβ-catenin compared to the wild-type NvAxin βcatBM-like sequence, and mutating both leucines to aspartic acid completely abolished interactions between NvAxin and Nvβ-catenin (Fig. 8C). In sum, these results suggest that the bilaterian βcatBM evolved as a sequence on the Axin-RGS with poor ability to interact with β-catenin and during pre-bilaterian evolution this sequence acquired a critical leucine sequence that permitted weak interactions with β-catenin.

**Figure 8.**
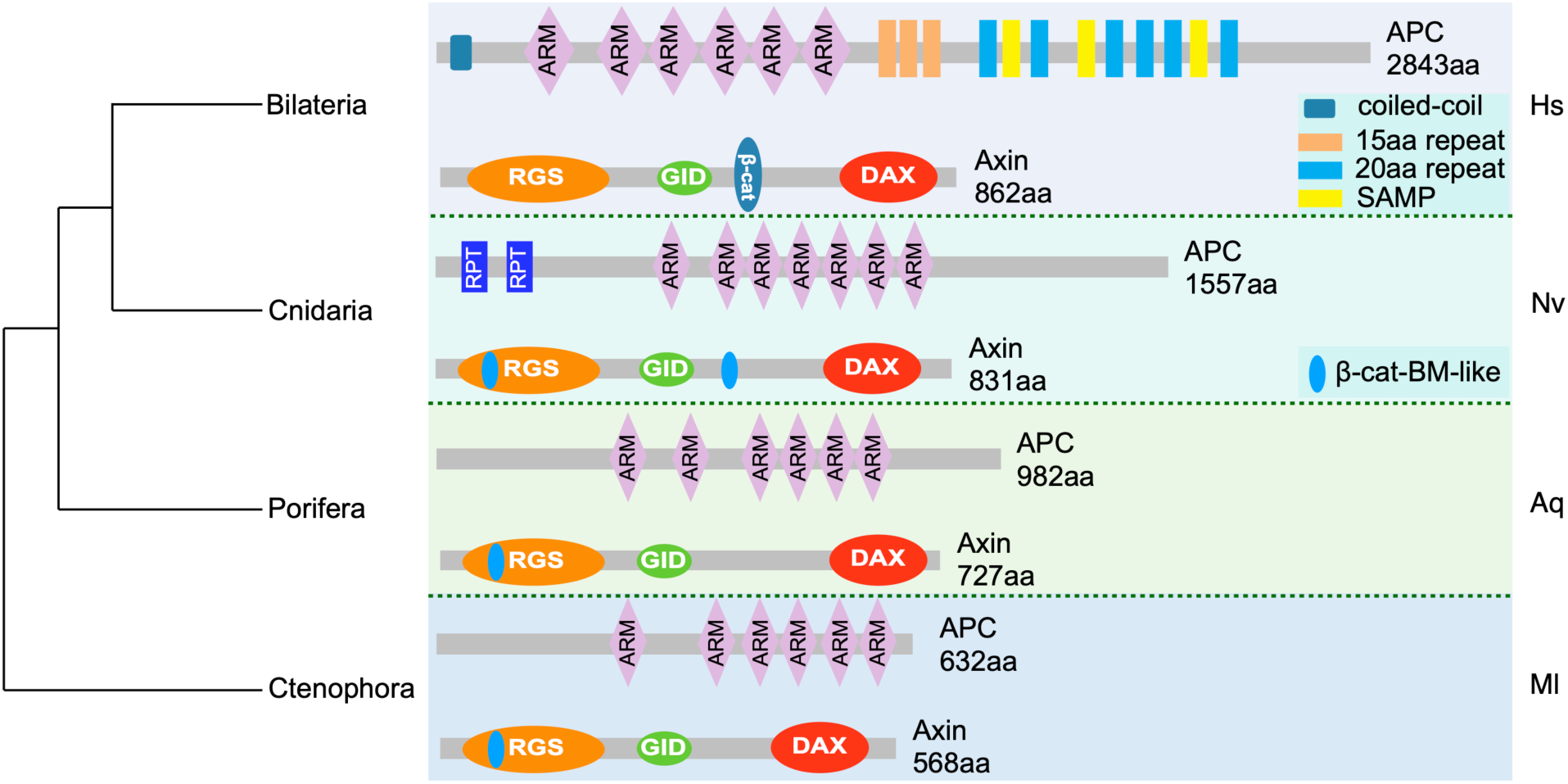
A model for the co-evolution of the β-catenin destruction complex scaffolding proteins Axin and APC-like/APC. The last common ancestor (LCA) to metazoans most likely had an Axin protein that lacked the canonical β-catenin-binding motif (βcatBM) but had a βcatBM-like motif embedded within the RGS domain with possibly promiscuous β-catenin-binding activity. The APC-like protein in this metazoan LCA had ARM domains with strong similarity to APC proteins in extant species. We do not have any evidence that the Axin-APC-like proteins in the metazoan LCA scaffolded a functional β-catenin destruction complex. There were relatively few changes in the main domains in Axin and APC-like in the LCA to the Porifera and later branching metazoans, but there was the acquisition of a critical leucine in the βcatBM-like motif embedded within the RGS domain that may have imparted a higher β-catenin-binding affinity to this motif. Based on the available evidence, we propose that in the Cnidarian-Bilaterian LCA, the βcatBM-like motif embedded within the RGS domain was duplicated with the duplicated copy inserted between the GID and DAX domains of Axin. The Axin and APC-like proteins in this LCA had several protein-protein interactions not reported in bilaterians and as such, may be weak interactions not previously defined in bilaterians that scaffolded a nascent destruction complex. In the Bilateria, the βcatBM on Axin evolved with the acquisition of additional critical amino acids including a critical histidine and the βcatBM-like motif in the RGS domain was lost, possibly through drift. The Axin protein in the Bilateria did not gain additional major domains, but the APC protein gained multiple domains that allowed it to interact strongly with several components of the destruction complex and over 100 other proteins (Nelson and Nathke, 2013). Here, the domains on the bilaterian APC have been simplified for clarity but the additional domains can be seen in Fig. 1D. Model was created using BioRender.

## Discussion

Studies on the evolution of metazoan signaling transduction pathways have been impeded by the limited understanding of their roles in early emerging taxa and the challenges to doing functional studies in these animals. We took advantage of the ability to experimentally manipulate cWnt signaling in *Nematostella* embryos and the *in vitro* assays we developed to identify direct protein interactions, coupled with the use of AI tools to study the evolution of the cWnt destruction complex. We show that *Nematostella* Axin and APC—which lack many of the domains required for bilaterian DC function—can effectively regulate the stability of cytoplasmic β-catenin using protein-protein interaction interfaces that to the best of our knowledge were previously undefined in bilaterians. We also traced the evolution of the critical bilaterian Axin β-catBM and provide evidence that it first appeared as a short β-catBM-like sequence embedded in the Axin-RGS domain in the earliest branching metazoans. Our analysis further indicates that during pre-bilaterian evolution the Axin β-catBM gained a leucine residue that permitted weak direct interactions between Axin and β-catenin. The β-catBM evolved in bilaterians through duplication of the β-catBM-like sequence in the cnidarian-bilaterian last common ancestor (LCA) with the duplicated sequence inserted towards the C-terminus of Axin. Our observations also suggest that in the Bilaterian lineage the duplicated β-catBM-like sequence gained a higher affinity for β-catenin through acquisition of additional amino acids, including a highly conserved histidine, and the original β-catBM-like sequence was lost, most likely through drift. Overall, this work has provided novel insight into the functional evolution of a major metazoan signal transduction pathway.

### NvAxin and NvAPC-like can assemble a functional destruction complex using novel protein binding interfaces

Axin and APC-like in cnidarians and other non-bilaterians lack the domains defined in bilaterians for assembling a functional cWnt DC. Hence, to investigate if the *Nematostella* DC could regulate cWnt signaling we carried out a series of experiments using readouts for cWnt signaling that we have previously established (Wikramanayake et al. 2003; Lee et al. 2007; Kumburegama et al. 2011; Röttinger et al. 2012; Sun et al. 2021). Our results showed that both NvAxin and NvAPC play a role in cWnt signaling, and moreover, we showed that NvAxin can also regulate cWnt signaling in sea urchin embryos (Fig. 4). We also used an *in vitro* protein-protein interaction assay to determine if NvAxin and NvAPC are able to directly interact with each other and with NvGSK-3β and Nvβ-catenin. This assay permitted us to identify previously undefined interactions between DC components. For example, our *in vitro* experiments showed that as predicted from NvAxin structure, this protein binds robustly to NvGSK-3β. This assay also allowed us to detect weak interactions between NvAxin and Nvβ-catenin that were unexpected because we were not able to identify a primary sequence that resembled the bilaterian β-catBM on NvAxin using SMART software. But using AI tools we were able to identify two β-catBM-like sequences on NvAxin. The *in vitro* assay also allowed us to show that NvAPC-like binds robustly to Nvβ-catenin and this interaction was also unexpected since NvAPC-like lacks the multiple domains that have been shown to bind β-catenin in bilaterians (Adamska et al. 2010; Roberts et al. 2011; Nelson and Näthke 2013; Pronobis et al. 2017; Abbott and Näthke 2023) (Fig. 1). Moreover, our *in vitro* experiments showed that NvAxin and NvAPC-like can interact directly even in the absence of the SAMP domains in NvAPC-like. Previous studies in *Drosophila* have shown that DmAPC2 can interact with Axin in the absence of the SAMP domains suggesting that some bilaterian APC proteins may also contain protein interaction interfaces that have not been precisely defined previously (Roberts et al. 2011). We suggest that in *Nematostella*, the interaction between NvAxin and NvAPC-like *in vivo* would likely bring β-catenin and GSK-3β together for phosphorylation of the β-catenin degron targeting the protein for degradation.

Other structural features of NvAPC-like indicated additional differences between the *Nematostella* and bilaterian cWnt DCs in the molecular mechanisms targeting β-catenin to the proteasome. Previous work has shown a clear role for NvAPC-like in regulating cWnt signaling in *Nematostella* (Kraus et al. 2016) and our results corroborated that work. However, the structure of NvAPC-like predicts that this protein would be unable to target β-catenin to the proteasome. For example, APC proteins from *Drosophila* and vertebrates have several 15 and 20 amino acid repeats that can bind to β-catenin and in addition, these proteins have 2-3 SAMP domains that are involved in dimerizing with Axin (Polakis 1995; Yang et al. 2006; Roberts et al. 2011; Nelson and Näthke 2013; Pronobis et al. 2015; Abbott and Näthke 2023) (Fig. 1D). The second 20aa repeat and an adjacent sequence are particularly significant because current models propose that these domains are critical for transferring phosphorylated β-catenin from Axin to APC prior to being moved to the proteasome for degradation (Kimelman and Xu 2006; Roberts et al. 2011; Pronobis et al. 2015). Strikingly, the NvAPC-like protein lacks any detectable 15aa and 20aa repeats or the SAMP domains but has seven predicted ARM repeats and two poorly characterized repeat regions (Fig. 1D). Hence, it appears that NvAPC-like resembles a severely truncated bilaterian APC protein that would likely be oncogenic in humans, and one that would not be predicted to regulate β-catenin degradation (Polakis 1995; Kinzler and Vogelstein 1996; Fodde et al. 2001; Schneikert and Behrens 2007). Despite these apparent structural deficits in cnidarian DC scaffolding proteins experimental data in a number of cnidarian species has shown that they can tightly regulate β-catenin protein stability in embryos. For example, when a β-catenin-GFP fusion protein is expressed by mRNA injection in *Nematostella* and other cnidarian embryos the protein is initially broadly expressed, but it is then cleared from the vegetal pole blastomeres of the embryos and very tightly restricted to nuclei of the animal pole blastomeres (Wikramanayake et al. 2003; Lee et al. 2007; Momose and Houliston 2007; Momose et al. 2008; Kumburegama et al. 2011; Röttinger et al. 2012). Hence, our observations raise important questions as to whether NvAPC-like has a role in shuttling phosphorylated β-catenin to the proteasome, or if cnidarians have a different mechanism to carry out this critical function. These are important questions that need to addressed in future studies.

Another striking observation from our phylogenetic analysis of the Axin and APC-like/APC proteins is the differential acquisition of domains between these proteins during evolution. While we have not studied Axin domain evolution beyond the four main domains (RGS, GID, βcatBM and DAX), Axin does not seem to have gained additional domains other than the βcatBM with a high-affinity for β-catenin. In contrast, the APC-like/APC proteins seem to have acquired many additional domains in the bilaterian lineage likely through gene fusion (see Fig. 1) (Adamska et al. 2010; Nelson and Näthke 2013; Abbott and Näthke 2023). This is particularly striking since Axin and APC-like/APC most likely co-evolved in the cWnt DC and it would be a reasonable prediction that changes in one protein would likely lead to changes in its partner protein. Some of the additional domains acquired by bilaterian APC are likely domains that are required for other functions of APC through its interactions with over 100 other previously identified proteins (Nelson and Näthke 2013). It is possible that modeling the co-evolution of Axin and APC-like/APC may lead to some insights into the evolutionary forces constraining the size of Axin while allowing the dramatic increase in domain complexity and size in bilaterian APC-like/APC.

### Reconstructing the evolution of the bilaterian Axin β-catenin binding motif

Our experimental work combined with the use of AI-enabled AlphaFold software allowed us to propose a plausible mechanism for the evolution of the bilaterian Axin βcatBM from pre-bilaterian antecedents. Insights into how the Axin βcatBM evolved came from AlphaFold’s identification of the NvAxin sequences underlying the weak but consistent interactions we noticed between NvAxin and Nvβ-catenin *in vitro* and *in vivo.* AlphaFold software predicted that two βcatBM-like sequences on NvAxin with similarity to the conserved bilaterian βcatBM could potentially interact weakly with Nvβ-catenin. But surprisingly one sequence was embedded within the NvAxin-RGS domain and the other was closer to the C-terminus of Axin where the βcatBM is found in bilaterian Axin (Fig. 8; Supplementary Fig. 8). Using AlphaFold we found a βcatBM-like sequence embedded in the Axin-RGS domains in a placozoan, a sponge and three ctenophores but we could not identify a second similar sequence closer to the C-terminus in these taxa as we did in Axin from *Nematostella* and several other cnidarians (Fig. 8; Supplementary Fig. 8,9). In addition, we could not identify a βcatBM-like sequence in any bilaterian Axin-RGS. Within the two βcatBM-like sequences in NvAxin and the single βcatBM-like sequence in TaAxin-RGS and AqAxin-RGS, there is a conserved leucine residue, also seen in the bilaterian Axin βcatBM, that has been shown to be critical for interaction between Axin and β-catenin in *Drosophila* (Kremer et al. 2010) (Fig. 8; Supplementary Fig. 8C,D,F). But the βcatBM-like sequences of NvAxin, TaAxin and AqAxin lack a highly conserved histidine that has been shown to be critical for Axin-β-catenin interactions in *Drosophila* (Kremer et al. 2010) (Supplementary Fig. 8C,D,F). Our analysis showed that the βcatBM-like sequence in the *Mnemiopsis* Axin-RGS lacked both the conserved leucine and histidine residues required for interaction of the *Drosophila* βcatBM with β-catenin and we showed that MlAxin and Mlβ-catenin could not directly interact *in vitro* (Supplementary Fig. 8E,F).

It is now established that ctenophores are likely the earliest emerging metazoan taxon (Dunn et al. 2008; Schultz et al. 2023). Hence, based on metazoan phylogeny we can infer that the putative ancestral βcatBM-like sequence that was present in the Axin-RGS domain in the “urmetazoan” acquired a leucine residue in the βcatBM-like sequence prior to the split of the poriferans from a last common ancestor (LCA) to placozoans, cnidarians and bilaterians. An alternative scenario is that the leucine was present in the Axin βcatBM-like urmetazoan and was lost in ctenophore Axin. Subsequently, after splitting from an LCA with cnidarians, bilaterian Axin acquired the critical histidine residue, and perhaps others, in the βcatBM-like sequence which provided an increased affinity between Axin and β-catenin. These data help reveal the molecular changes that occurred prior to the stabilization of Wnt/β-catenin regulation that appears in the majority of bilaterian clades.

### Evolution of the bilaterian cWnt destruction complex

The cWnt pathway is one of only two signal transduction pathways regulating development that are restricted to all metazoans (Richards and Degnan 2009; Babonis and Martindale 2017; Richter et al. 2018) and it plays diverse roles in cellular and developmental processes and in homeostasis (Liu et al. 2002; MacDonald et al. 2009; Petersen and Reddien 2009; Loh et al. 2016). One highly conserved function of the cWnt pathway is its role in establishing the first embryonic axis in bilaterians (anterior-posterior) and cnidarians (oral-aboral) (Wikramanayake et al. 2003; Petersen and Reddien 2009; Loh et al. 2016; Holstein 2022). We and others have also shown that the cWnt pathway also plays a critical role in segregating germ layers in cnidarians and invertebrate bilaterians (Wikramanayake et al. 1998; Logan et al. 1999; Imai et al. 2000; Wikramanayake et al. 2003; Henry et al. 2008; Darras et al. 2011; McCauley et al. 2015; Holstein 2022). Considering the conservation of the role of cWnt signaling in these fundamental processes it is remarkable how divergent the regulation of the cWnt destruction complex appears to be between early branching metazoans and Bilateria. Below, we suggest a model for the evolution of the cWnt DC that argues that the cWnt DC in *Nematostella* is not “divergent” but it resembles the ancestral cWnt DC in the cnidarian-bilaterian LCA that was functionally sufficient to regulate cWnt signaling in animals that had relatively simple body plans. But this ancestral DC had promiscuous interactions that increased the evolvability of the cWnt pathway during its broad deployment as bilaterians underwent a massive radiation to evolve taxa with complex body plans.

We know very little about cWnt signaling in ctenophores (Pang et al. 2010; Walters et al. 2025) and it has also been difficult to study embryonic development in two other early emerging taxa, poriferans and placozoans, due to the challenges in accessing their embryos. Hence, our current focus has been on reconstructing the ancestral cWnt DC in the cnidarian-bilaterian LCA, since we can experimental manipulate *Nematostella* development and there is accumulated knowledge on the developmental roles of the cWnt pathway in cnidarian development. We propose that the ancestral DC that was present in the LCA of Bilateria and Cnidaria was scaffolded by an APC-like protein that bound β-catenin and Axin, and an Axin protein that bound GSK-3β and had two βcatBM-like sites that had low-affinity β-catenin-binding activity. This DC was capable of regulating β-catenin stability in the cWnt pathway in the cnidarian-bilaterian LCA. Whether this DC was able to directly target β-catenin for degradation through the proteasome is not known at this time but regardless of the mechanism, it had to be very different from how β-catenin is targeted for degradation in the bilaterian cWnt DC (Roberts et al. 2011; Ranes et al. 2021).

Following the split of the bilaterians from the cnidarian-bilaterian LCA, the βcatBM-like site at the C-terminus positioned between the GID and DAX domains of bilaterian Axin gained amino acids that increased its affinity for β-catenin possibly by gaining the functionally significant histidine residue seen in the bilaterian βcatBM (Kremer et al. 2010). Likely in parallel, APC-like gained a long carboxy-terminal region that led to the acquisition of redundant β-catenin and Axin-binding sites. In addition, some bilaterian APC proteins gained domains that are involved in processes such as oligomerization and interaction with microtubules (Nelson and Näthke 2013; Abbott and Näthke 2023) (Fig. 8). The mechanism for domain addition to APC is unknown but it is likely due to gene fusion since that is considered to be the predominant mechanism for domain evolution in eukaryotic proteins and it generally occurs at the N- or C-termini of proteins (Patthy 1999; Liu et al. 2005; Weiner et al. 2006).

We were unable to identify a βcatBM-like sequence in the Axin-RGS domain of all the bilaterians we examined (listed in Table 1). This sequence was most likely lost through drift, since with the acquisition of APC domains that interacted with high affinity with the Axin RGS domain of the βcatBM-like sequence embedded in the RGS would not have been able to interact with β-catenin if it had an earlier function in binding β-catenin. The overall consequence of these changes may have been the evolution of higher-affinity protein-protein interactions within the DC that allowed a more efficient capture of free β-catenin from the cytoplasm and an increased level of redundancy for β-catenin sequestration that served to buffer this critical regulatory module from misregulation (Peterson-Nedry et al. 2008; Roberts et al. 2011) (Fig. 8C). This idea is consistent with modeling studies which have shown that signaling pathways evolve increased levels of complexity under standard evolutionary conditions, and this complexity also provides increased robustness (Soyer and Bonhoeffer 2006). It is also likely that the increased number of interfaces provided by the bilaterian Axin and APC proteins provided additional targets for context-dependent regulation of the cWnt DC during the evolution of complex body plans in the Bilateria. In conclusion, we suggest that the cnidarian-bilaterian LCA cWnt DC had promiscuous protein interactions that may have increased the evolvability of the cWnt pathway during its broad deployment during bilaterian development and evolution. We believe that the approach we have taken in this study illustrate the power of leveraging biodiversity to gain insight into the evolution of protein function and protein-protein interactions networks that regulate evolution and development.

## Materials and Methods

### Animal care, spawning and culture

Adult *N*. *vectensis* were spawned and eggs prepared for microinjections as previously described (Wijesena et al. 2022).

Adult *Lytechinus variegatus* (Lv) were obtained from Duke University Marine Lab (Beaufort, NC) or Pelagic Corporation (Sugarloaf Key, FL). Spawning of animals and collection and treatments of gametes was done as previously described (Byrum et al. 2009; Sun et al. 2021).

### Cloning full length NvAxin and Reverse Transcriptase-PCR analysis

Full length *N. vectensis* Axin (NvAxin) cDNA was PCR-amplified from egg stage using primers designed against the NvAxin sequence (NEMVEDRAFT_v1g182113). The primer sequences were: NvAxin FL forward primer: CGCGCGGATCCATGAGCGAAGAGTTGTGTAG, and NvAxin FL reverse primer: CGACCATCGATTGAGTCTACTTTTTCTACTTTGC. Egg, cleavage, blastula, gastrula and planula stage embryos were collected separately for RNA extraction using Trizol reagent (cat# 15596026, Invitrogen). Approximately 1 µg of total RNA was used for cDNA synthesis using SuperScript IV first strand synthesis system (cat# 18091200, Invitrogen). For PCR, NvAxin cDNA was amplified from different stage embryos. The primers used to amplify NvAxin for RT-PCR were: NvAxin FP: ATGAGCGAAGAGTTGTGTAG, and NvAxin RP: CGTGCATCTTGCTCGTCGTG.

The PCR procedures followed the protocol provided with the Q5 high fidelity DNA polymerase kit (cat# M0491, NEB) with thermal cycling conditions, 98°C for 30 seconds, 35 cycles with (98°C 10 seconds, 62°C 30 seconds, 72°C 20-30 seconds/kb) and 72°C for 2 minutes. The wildtype NvAxin was inserted into the GFP-pCS2^+^ vector by BamH1 and Cla1 restriction enzyme digestion and ligation with T4 DNA ligase.

### Making NvAxin domain deletion constructs by site-directed mutagenesis

NvAxinΔRGS, NvAxinΔGID, NvAxinΔDAX constructs were made using the Q5 Site-directed mutagenesis kit (E0554, NEB). Primers were designed to delete the RGS, GID and DAX domains separately. The following primer sequences were used: NvAxin ΔRGS forward primer, CAGGCTCACAGAGACACC, and NvAxin ΔRGS reverse primer, GAGTTAGCCCATCGCTCG; NvAxin ΔGID forward primer, TGAGAGTGAATATGGCCAG, and NvAxin ΔGID reverse primer, GGATTTTTGGCTTCCTTTG; NvAxinΔDAX forward primer, CAATCGATTCGAATTCAAG, and NvAxinΔDAX reverse primer, TTGTGTTTCTCCTTTTTCC. To perform the PCR procedure, the NvAxin-GFP pCS2+ plasmid was mixed with pairs of primers, the Q5 hot start high-fidelity 2X master mix and nuclease free water to perform PCR procedure, followed by kinase, ligase & Dpnl (KLD) treatment.

### Microinjection of morpholino antisense oligonucleotides and mRNAs

For knockdown experiments, morpholino antisense oligonucleotides to Axin or APC-like (NvAxin MO or NvAPC MO) and a standard control morpholino (Control MO) to a random nucleotide sequence (5’-CCTCTTACCTCAGTTACAATTTATA-3’) were synthesized at Gene Tools, LLC (Eugene, OR). The NvAxin MO: TACACAACTCTTCGCTCATCTCACG or NvAPC-like MO: ATCCTCTTCAAAATCGTCCATGTTG were designed to span the start codon of the Axin or APC-like transcripts. Each MO was injected at a final concentration of 2 mM in 30-40% glycerol and dextran rhodamine 0.2 µg/µl (D1824, Invitrogen).

For overexpression experiments in *N. vectensis* or *L. variegatus*, the NvAxin-GFP, NvAxin ΔRGS-GFP, NvAxin ΔGID-GFP, NvAxin ΔDIX-GFP, SpAxin-GFP, Nvβ- catenin-mCherry, and Spβ-catenin-mCherry, NvAxin GID-GFP, and a control GFP mRNA were all cloned into the pCS2+ vector. The pCS2+ vectors containing the respective cDNAs were linearized with NotI and mRNA was transcribed using the SP6 mMessage mMachine Kit (Ambion® ThermoFisher Scientific). Dominant negative NvTcf (NvdnTCF) (Röttinger et al. 2012) was used for downregulating cWNT signaling. This construct was in the Venus-pDEST expression vector. The dnTCF-Venus construct was linearized with Acc65I and *in vitro* transcribed as previously described (Röttinger et al. 2012). The mRNAs were mixed in 40% glycerol to a final concentration of 0.5 µg/µl. *GFP* mRNA was overexpressed as a control at the same molar ratio as previously described (Bince and Wikramanayake 2008).

For co-overexpression experiments in *N. vectensis* or *L. variegatus*, *NvAxin-GFP*, *NvAxin ΔGID-GFP*, *SpAxin-GFP*, *Nvβ-catenin-mCherry*, *Spβ-catenin-mCherry* mRNAs were injected at a final concentration of 0.5 µg/µl, 0.5 µg/µl, 0.5 µg/µl, 0.2 µg/µl, 0.2 µg/µl respectively in 40% glycerol.

For each experiment, more than 100 embryos were injected per MO or per mRNA. The experiments were repeated at least three times for biological replication (Sun et al. 2021; Wijesena et al. 2022).

### CRISPR/Cas9 mediated knockouts and microinjections

To knock out *NvAxin* or *NvAPC-lik*e, the synthetic guide RNAs (sgRNA; Synthego, Inc.) and Cas9 complex (RNP complex) were introduced into *N. vectensis* embryos by microinjection into zygotes. The *NvAxin* or *NvAPC-like* single guide RNAs were designed using CRISPRscan and CHOPCHOP as previously described (Salinas-Saavedra et al. 2018). Each single guide RNA was resuspended to 50 µM and stored at −80°C. Cas9 was obtained from PNA bio (Cat.# CP01). Cas9 was reconstituted to 5 mM and was aliquoted for single use and stored at −80°C.

A mixture (2.5 µl) containing sgRNA (12 µM), Cas9 (0.6mM), 40% Glycerol, dextran rhodamine (0.2 µg/µl) and nuclease free H_2_O was prepared and 1 µl of mixture was loaded into needles for microinjection. The same amount of Cas9 only was injected as a control. *NvAxin* sgRNA was used to introduce mutations into *NvAxin*, and *NvAPC-like* sgRNA was used to introduce mutations into *NvAPC-like*, and the mixture of *NvAxin* and *NvAPC-like* sgRNAs were used to mutate both genes in single embryos.

### Genomic DNA extraction from single embryos and validation of DNA editing by CRISPR/Cas9

Single *NvAxin* gRNA injected embryos were collected into 0.2 ml PCR tubes and lysed by adding 20 µl of extraction buffer for 100 µl buffer: 1 µl 1M Tris, pH8.0; 0.3 µl Tween 20; 0.2 µl 0.5M EDTA; 5 µl 1M KCl; 0.3 µl NP40; 2.5 µl proteinase K (20 mg/ml); nuclease free water). The tubes with lysed embryos were incubated for 3 hr at 55°C in a thermal cycler with periodic vortexing and the reaction was inactivated by heating the tubes at 98°C for 5min. For the PCR analysis of *NvAxin* gRNA injected embryos, 5 µl of genomic DNA was used as a template to carry out the PCR reaction as described above for NvAxin cloning and then the amplicon was gel extracted, purified and sequenced at Eurofins Scientific. The following primers were used to generate the NvAxin amplicons from genomic DNA from embryos edited with the *NvAxin* gRNA: forward primer: ATGGCCAGTCTAAAAAGTCTATCC, and reverse primer: CTGGTGGTTCAATGTCAAGAACGGTAT.

### Quantitative PCR (qPCR)

To evaluate the effects of injected gRNAs, MO or mRNA on lineage-specific gene expression, quantitative PCR (qPCR) was performed on *N. vectensis*, or *L. variegatus* cDNA samples synthesized from total RNA that was extracted from gRNA-, morpholino-or mRNA-injected embryos. The control and treated samples were collected at the blastula stage; *N. vectensis* embryos were at 24 hpf at 17°C and *L. variegatus* embryos were at 6 hpf at 25°C. Total RNA was isolated using the Trizol reagent solution (cat# AM9738, Invitrogen). The cDNA was synthesized from 50 -100 ng total RNA of control and experimental embryos respectively using the qScript cDNA Synthesis kit (Quanta Biosciences, Beverly). The PerfeCTa SYBR Green FastMix (Quanta Biosciences, Beverly) was used for assembling the qPCR reactions. Each experiment was repeated 3 times with separate batches of embryos and each PCR reaction was done in triplicate.

The expression of selected genes was analyzed using the delta delta Ct (2^-ΔΔCt^) method and using GADPH as a control as previously described (Sun et al. 2021). Statistical significance was determined using the Unpaired t test without specific fold change cut off. The values represent the fold change relative to control. All the *L. variegatus* primer sequences were obtained from David McClay’s laboratory at Duke University.

### Whole mount in situ hybridization (WMISH)

*NvAxin* expression during early development was determined using whole mount in situ hybridization (WMISH). The *NvAxin* in situ hybridization probe was made using a pair of primers specifically targeting NvAxin: NvAxin-F: 5’ CGAAGGTAGAGCCGTATC 3’, and NvAxin-R:

ATTTAGGTGACACTATAGGTCCGTCAGAGCATCACT, and was designed to contain an Sp6 promoter for *in vitro* RNA synthesis. Embryo fixation and WMISH were performed as previously described (Salinas-Saavedra et al. 2018). Embryos were fixed in 4% paraformaldehyde with 0.2% glutaraldehyde in PTw (1XPBS+ 0.1%Tween-20) for 2 min at room temperature, and then fixed in 4% paraformaldehyde in PTw for 1 hr at 4 °C. In each experiment, a final concentration of 1 ng/µL anti-sense *NvAxin* probe was used which was hybridized at 63°C for 2 days. *NvAxin* expression was detected using NBT/BCIP as a substrate for the alkaline phosphatase conjugated to anti-digoxigenin fab fragments (1:5000; Roche).

### F-actin and nuclear staining

The morphology of *dnTcf-*, *NvAxin-*, *SpAxin-* and control *GFP*-RNA injected embryos was examined using embryos collected at late gastrula stage (48 hpf at room temperature) that were fixed and stained for F-actin and nuclear staining. Embryos were collected and fixed as previously described with modifications (Salinas-Saavedra et al. 2018). Briefly, embryos were first fixed in fix1 (100 mM HEPES pH 6.9; 0.05 M EGTA; 5 mM MgSO4; 200 mM NaCl; 1x PBS; 3.7% Formaldehyde; 0.2% Glutaraldehyde; 0.5% Triton X-100; and autoclaved ultrapure water) for 2-3 minutes and then fixed in fix2 (100 mM HEPES pH 6.9; 0.05 M EGTA; 5 mM MgSO4; 200 mM NaCl; 1x PBS; 3.7% Formaldehyde; 0.05% Glutaraldehyde; 0.5% Triton X-100; and autoclaved ultrapure water) for one hour at room temperature on the rocker. Fixed embryos were rinsed in PBT (X% Tween 20 in 1 x PBS, pH7.4) for 30 mins x 4 times. Embryos were blocked with 5% normal sheep in PBT buffer for 1 hour at room temperature on the rocker then rinsed with PBT for 30 mins x 4 times. For the F-actin staining, embryos were stained with Phalloidin in PBT for 1 hour and then rinsed with PBT for 10 min x 2 times. For nuclear staining, embryos were incubated in DAPI in PBT for 1 hour and then rinsed in 1xPBS (PH 7.4) for 10 min x 3 times. Following these washes, the embryos were dehydrated using a gradient of isopropanol (50, 75, 90, 100%, 100%). The embryos were then cleared with 1:2 benzyl benzoate: benzyl alcohol (BB:BA) for 1 min x 2 times and mounted in BB:BA for imaging using scanning confocal microscopy.

### In vitro Transcription and Translation (TNT) and Western blot analysis

All GFP, mCherry, Flag and their fused constructs were in vitro transcribed and translated using the TnT® Quick Coupled Transcription/Translation System with SP6 RNA polymerase promoter respectively (cat # L2080, Promega). Plasmids (at 1 µg each) containing the respective cDNAs coding for the various fusion proteins were incubated with TNT quick master mix at 30°C for 90 minutes. Protein expression was examined using Western blot analysis with anti-GFP (cat# sc9996), anti-mCherry antibodies (PA5-34974) or anti-flag (cat# 740001) respectively.

Protein-protein interaction was tested using ChromoTek GFP-Trap® Magnetic Agarose following instructions to perform immunoprecipitation (IP) assay (cat # gtma, Proteintech Group, INC., Rosemont, IL). Denatured IP protein samples were examined on an 8% SDS-PAGE gel followed by Western blot analysis. Western blot was done as previously described (Sun et al. 2024).

### Split luciferase assay

N- and C-terminal fragments of click beetle green (CBG) split luciferase, including polypeptide linkers, were amplified and blunt-end-ligated into pSPE3-RfA vectors in a single restriction-ligation reaction. Coding sequences for *N. vectensis* and S*. purpuratus* Axin and β-catenin were cloned into pENTR using D-TOPO cloning kit (Invitrogen K240020) and recombined into CBG-tagged destination vectors via Gateway cloning (Invitrogen 11791020). Nanoluciferase (NanoLuc) was similarly cloned and recombined into pSPE3-RfA.

Recombined Axin and β-catenin constructs were linearized, purified, and used for in vitro mRNA synthesis (Invitrogen AM1348). Equimolar CBG-tagged *Axin* and *β-catenin* mRNA mixtures, along with NanoLuc mRNA and fluorescent dextran, were microinjected into *Nematostella* zygotes. At 28 hours post-fertilization(hpf), live embryos were collected, and luminescence was measured after adding D-luciferin substrate (Goldbio LUCNA-100). Embryos were then lysed with ONE-Glo EX Luciferase Assay Reagent (Promega N1610), and further measurements were taken. NanoDLR Stop & Glo Reagent (Promega N1610) was used to stop the CBG reaction and initiate NanoLuc luminescence. NanoLuc signal was used to normalize protein levels and CBG intensities. Average NanoLuc signal from 15-20 minutes after the addition of substrate was used to determine ratios of protein levels. Average relative intensities across timepoints were calculated for each construct combination.

Primers:

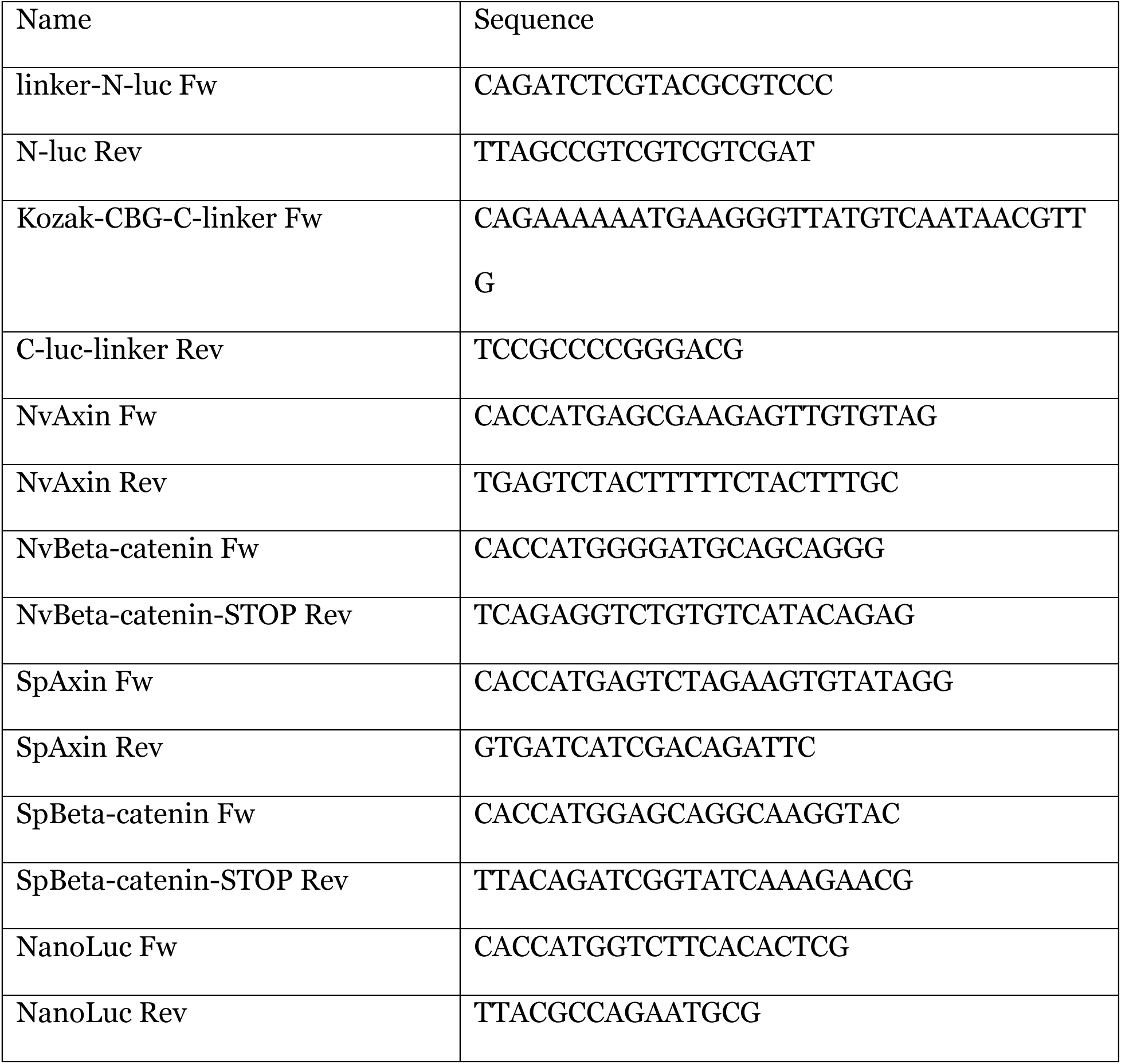

### AlphaFold 3 Protein-Protein interaction prediction

Computational methods were used to predict protein-protein interactions among the components of the β-catenin destruction complex in *S. purpuratus* (sea urchin) and *Nematostella*. Protein sequences for the individual components of the β-catenin destruction complex were used as input in AlphaFold 3. The prediction accuracy was assessed using the predicted align error (PAE) and confidence scores such as the predicted TM-score (PTM) and inter-chain predicted TM-score (iPTM), which are indicators of the reliability of the predicted interactions. A PTM score above 0.5 indicates that the overall predicted fold for the complex may resemble the true structure. iPTM Values greater than 0.8 represent confident, high-quality predictions, while values below 0.5 suggest likely failed predictions. ipTM scores between 0.5 and 0.8 fall into an intermediate zone where predictions may be correct or incorrect21. AlphaFold 3 allowed for the visualization of possible binding interfaces and the determination of whether specific regions, such as the β-catenin binding domains, could facilitate protein-protein interactions, even in cases where canonical domains were not evident.

## Acknowledgements and funding sources

This work was partially supported by NASA (80NSSC18K1067) to AHW and MQM and NSF (1755364) to MQM. We thank Ryan Christopher Vignogna, Lin Luan, and Le Li for help with AlphaFold analysis.

---

**Supplementary Table 1.** The primary protein interaction domains on Axin proteins from bilaterian and non-bilaterian taxa. In bilaterians (black), Axin proteins contain APC- (RGS), GSK-3β- (GID), β-catenin- (β-cat), and Disheveled-(DAX) binding domains. In non-bilaterians (red), the conserved β-catenin binding domain between the GID and DAX domains is absent.

| Axin Domains<br>Species Name & Accession # | RGS | GID | β-cat | DAX |
| --- | --- | --- | --- | --- |
| *Strongylocentrotus purpuratus* (XP_003729076) | ✓ | ✓ | ✓ | ✓ |
| *Saccoglossus kowalevskii* (NP_001158475) | ✓ | ✓ | ✓ | ✓ |
| *Platynereis dumerilii* (ANS60441) | ✓ | ✓ | ✓ | ✓ |
| *Crassostrea gigas* (EKC24785.1) | ✓ | ? | ✓ | ✓ |
| *Branchiostoma floridae* (XP_035695674) | ✓ | ✓ | ✓ | ✓ |
| *Homo sapiens* (NP_003493) | ✓ | ✓ | ✓ | ✓ |
| *Mus musculus* (AAC53285) | ✓ | ✓ | ✓ | ✓ |
| *Xenopus laevis* (NP_001081874) | ✓ | ✓ | ✓ | ✓ |
| *Danio rerio* (BAA92439) | ✓ | ✓ | ✓ | ✓ |
| *Hofstenia miamia* (KJ658749.1) | ? | ✓ | ✓ | ✓ |
| *Tribolium castaneum* (NP_001280495) | ✓ | ? | ✓ | ✓ |
| *Drosophila melanogaster* (NP_733336) | ✓ | ✓ | ✓ | ✓ |
| *Apis mellifera* (XP_006564444) | ✓ | ✓ | ✓ | ✓ |
| *Nematostella vectensis* (XP_048576223) | ✓ | ✓ | x | ✓ |
| *Mnemiopsis leidyi* (ML04208a) | ✓ | ✓ | x | ✓ |
| *Amphimedon queenslandica* (NP_001266244) | ✓ | ✓ | x | ✓ |
| *Trichoplax adhaerens* (XP_002109778) | ✓ | ✓ | x | ✓ |

**Supplementary Figure 1.**
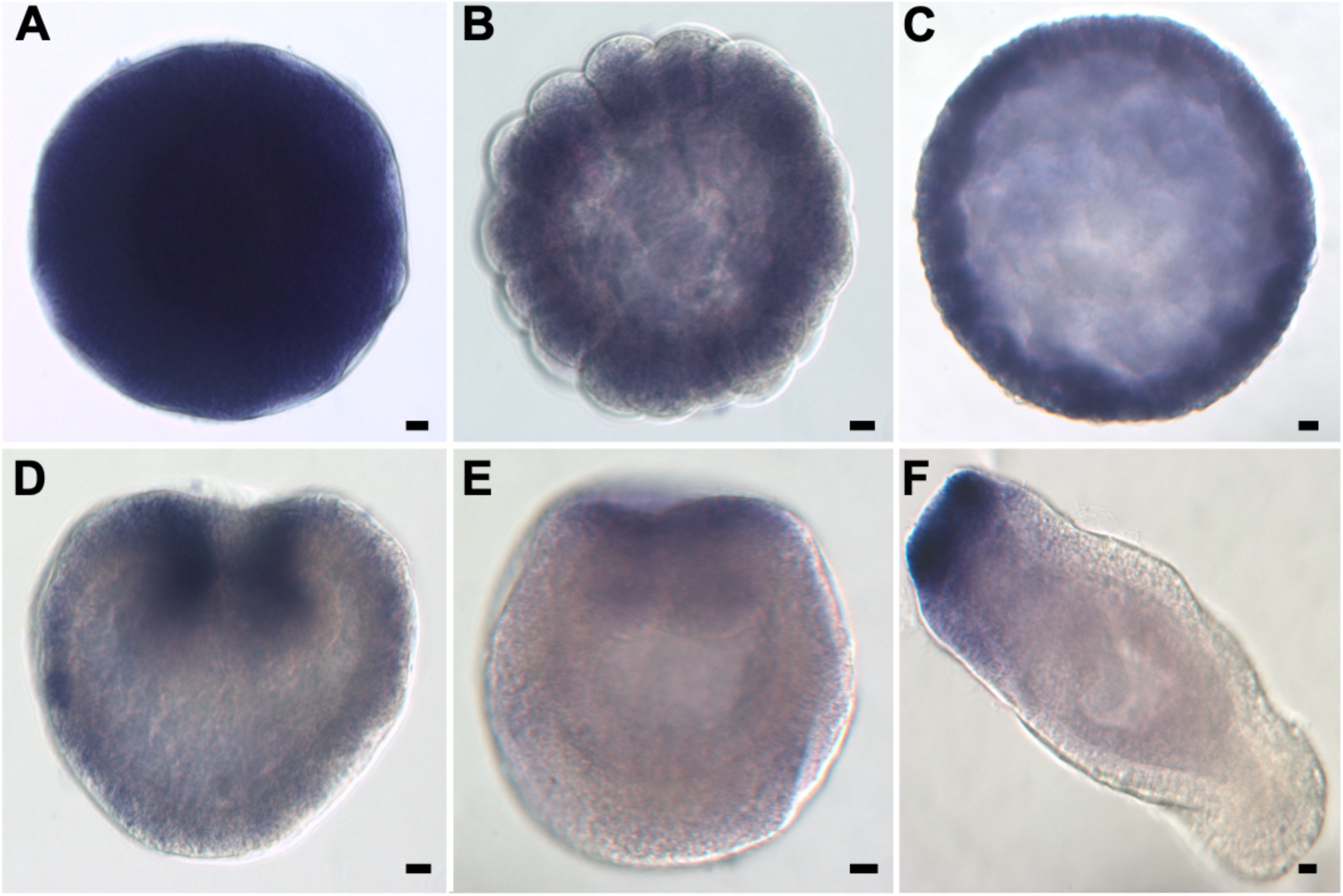
Whole mount RNA in situ hybridization shows the expression of *Axin* in *Nematostella* eggs and embryos. Axin is ubiquitously expressed in the egg, and during cleavage and blastula stages (A-C), however Axin is asymmetrically expressed at the animal pole at early and late gastrula stage (D, E) and the oral side at planula stage (F). Scale bar = 10 µm.

**Supplementary Figure 2.**
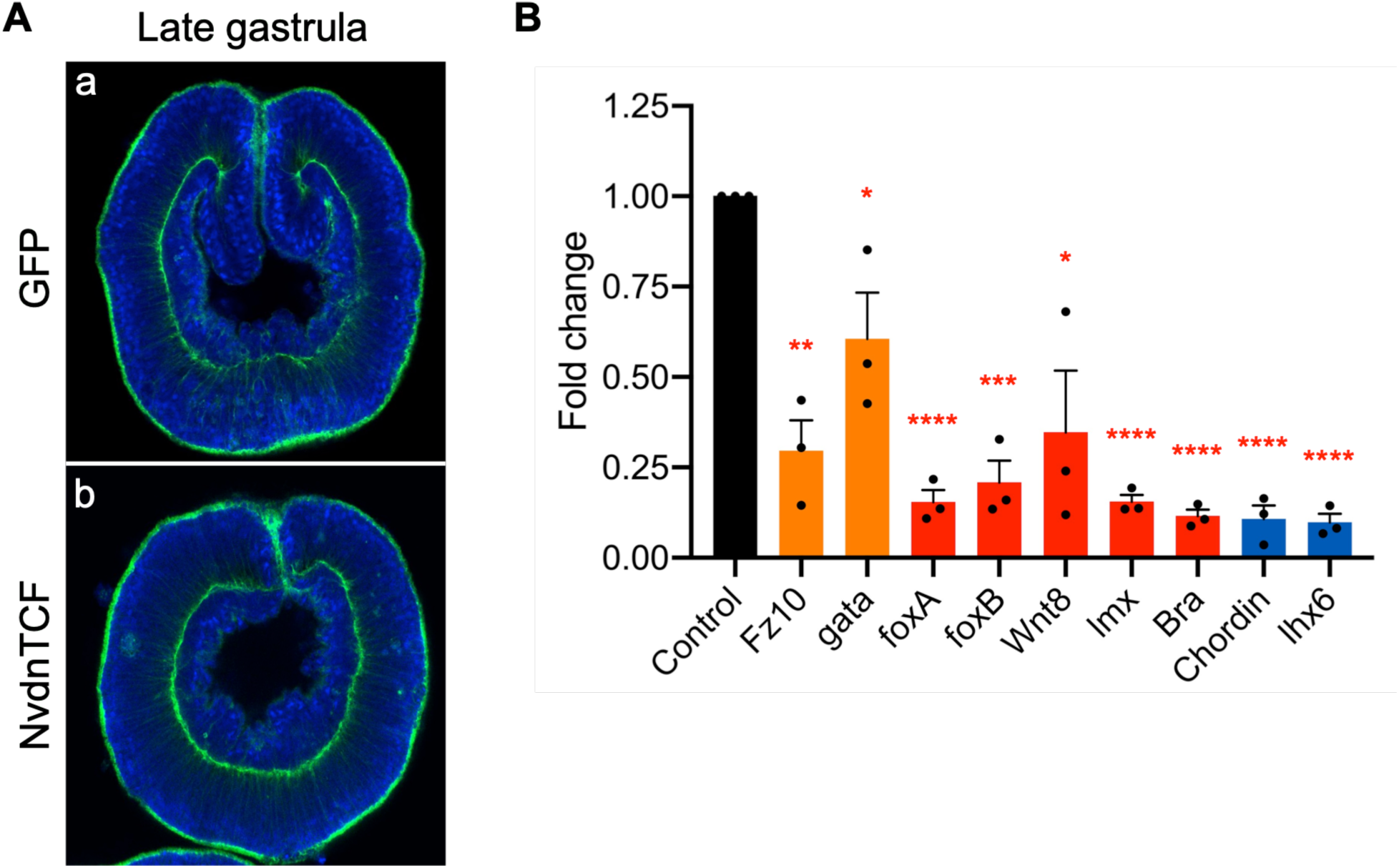
The effect of NvTcf knockdown on primary invagination and endomesodermal and pharyngeal gene expression in *Nematostella* embryos. (A) Compared to GFP overexpressing control embryos (a), NvdnTCF-overexpressing embryos (b) initiated primary invagination but failed to maintain a normal archenteron epithelium at the late gastrula stage. (B) qPCR analysis of NvdnTCF-overexpressing embryos at 24 hpf (hours post fertilization) showed that all genes expressed in cells derived from animal pole blastomeres (orange, red, and blue bars) are downregulated. qPCR experiments were replicated with three separate batches of embryos with three technical replicates in each experiment. GAPDH was used as the internal control. Statistical significance was determined using the Unpaired t test. ****, p<0.0001; ***, p<0.001; **, p<0.01; *, p<0.05; n=3.

**Supplementary Figure 3.**
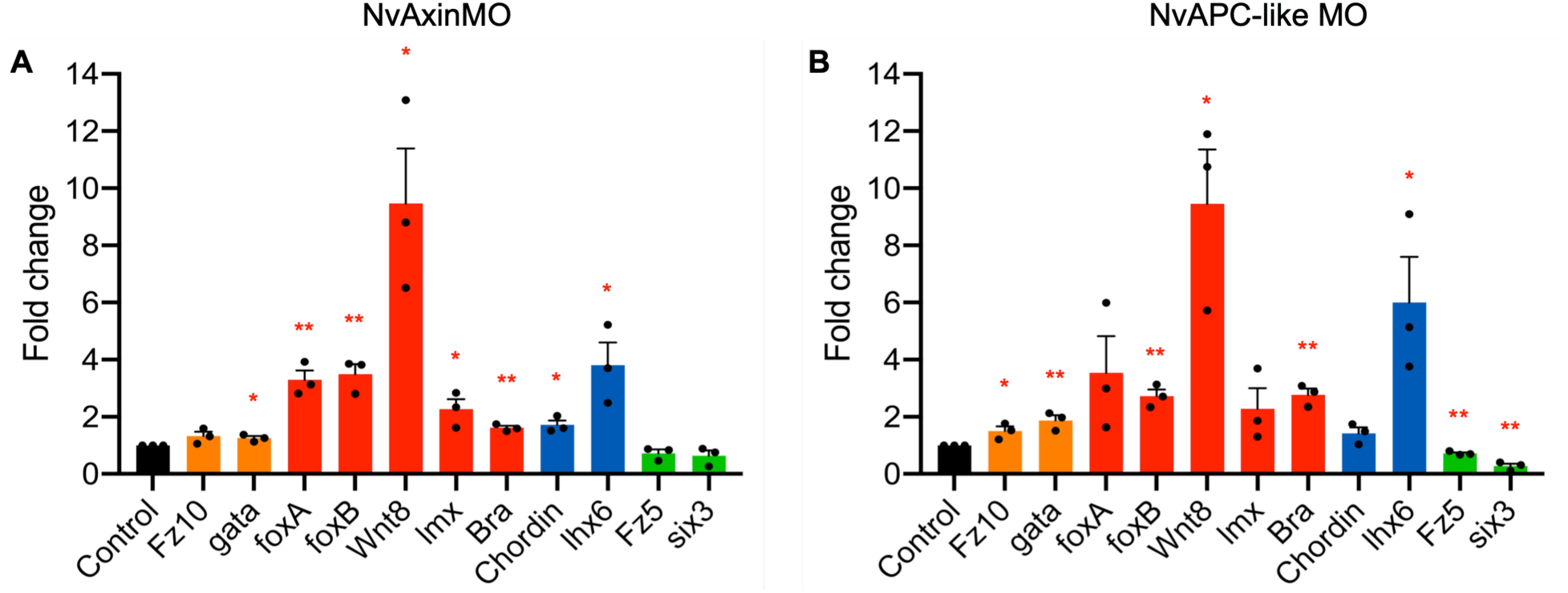
Morpholino mediated knockdown of Axin or APC-like upregulates endomesoderm gene expression in *Nematostella* embryos. (A,B) Endomesoderm and pharynx gene markers were upregulated in NvAxinMO or NvAPC-like MO-injected embryos compared to control MO (black bars). The qPCR experiments were replicated with three separate batches of embryos, with three technical replicates in each experiment. GAPDH was used as the internal control. Statistical significance was determined using the Unpaired t test. **, p<0.01; *, p<0.05; n=3.

**Supplementary Figure 4.**
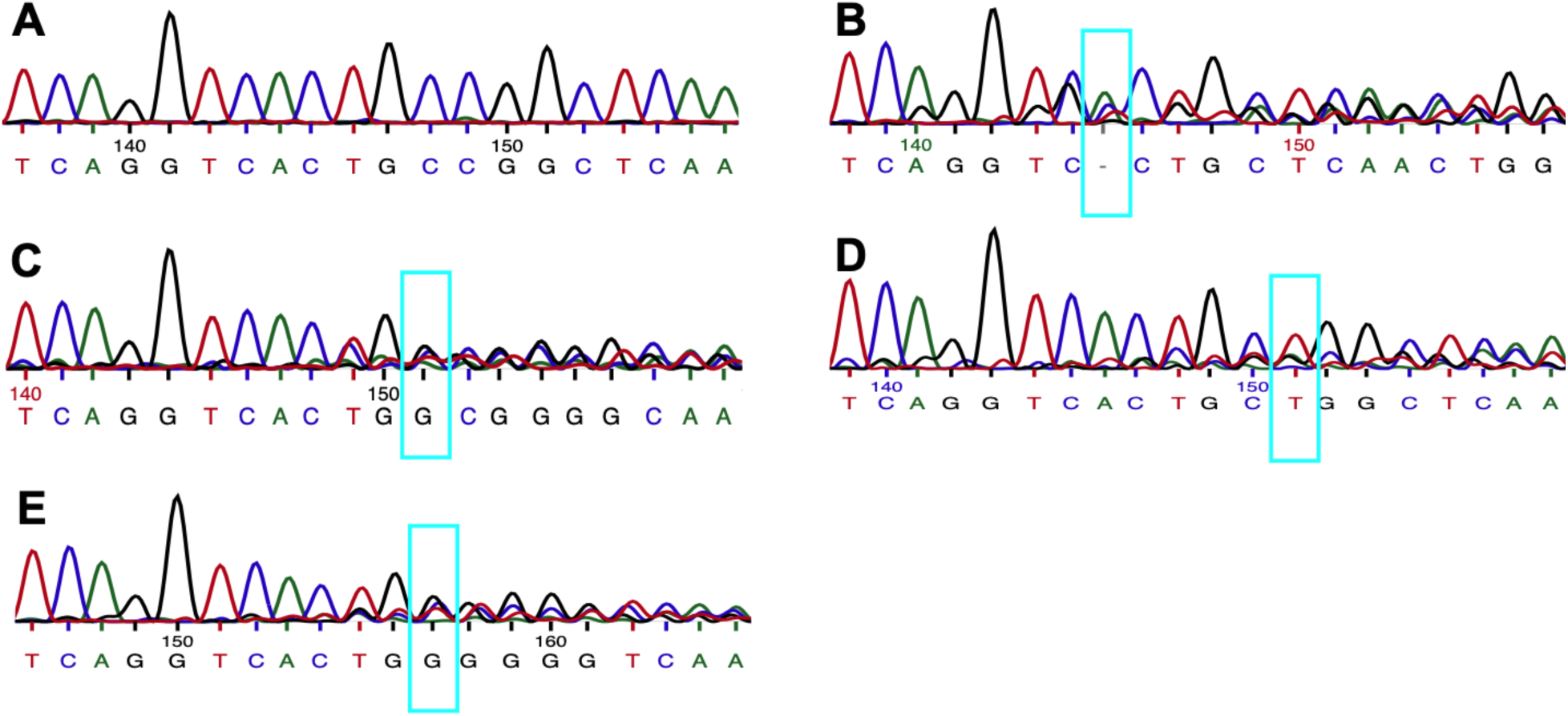
Confirmation of successful CRISPR/Cas9-mediated gene editing with Sanger sequencing using a single injected embryo from each group. Compared to the Cas9 only injected embryo (A), the sequencing chromatograms showed that injection of NvAxingRNA/Cas9 induced deletions or mutations (cyan box) at the targeted region of the Axin gene (B-E).

**Supplementary Figure 5.**
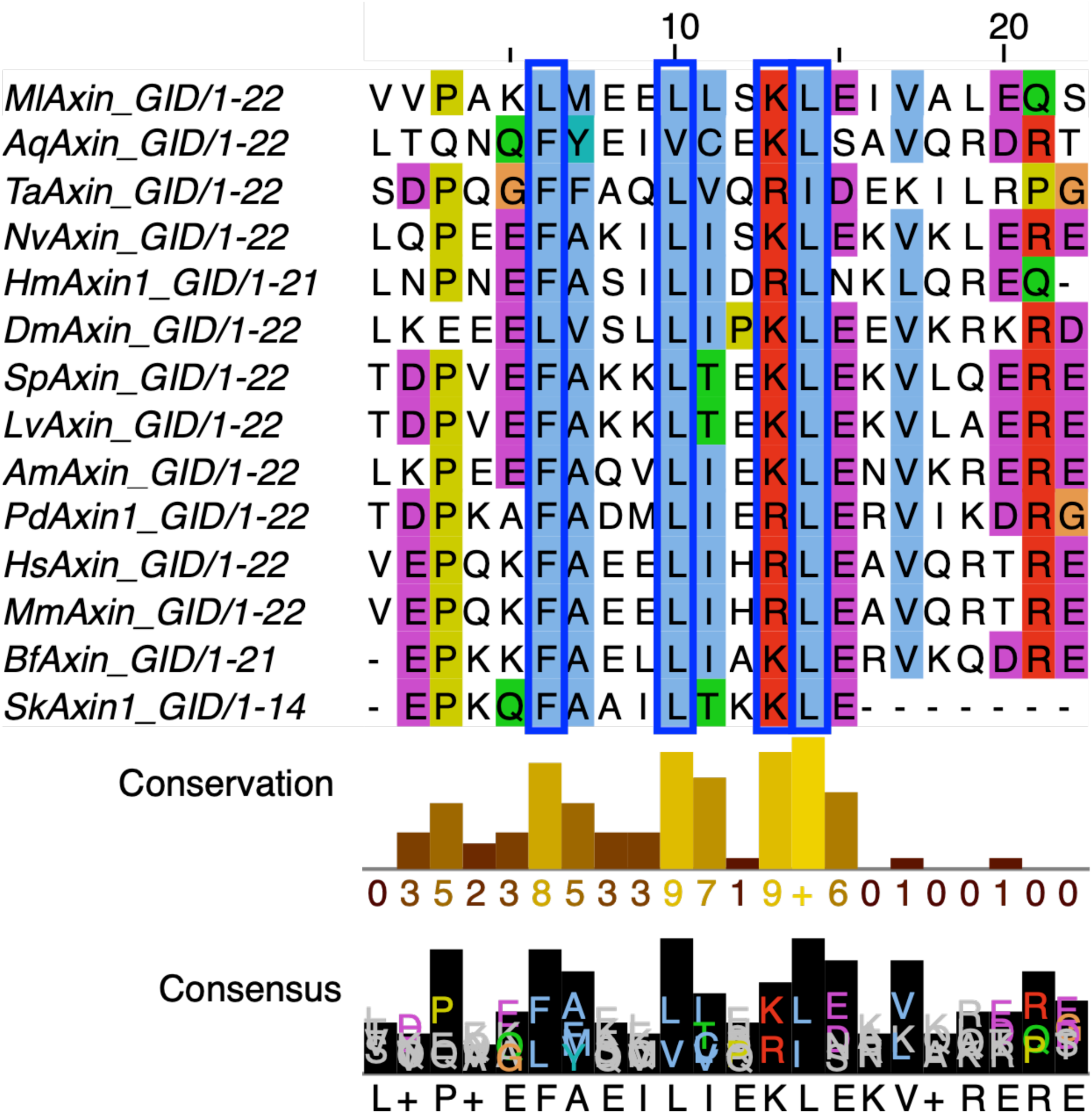
The protein sequence alignment of Axin GID domains in various metazoan taxa. The top panel shows the alignments of amino acids sequence using Clustal Omega and Jalview. Conserved amino acids are highlighted in different colors, while gaps are represented in white. The bottom panel is the analysis of conservation and consensus. *Ml*, *Mnemiopsis leidyi*; *Aq*, *Amphimedon queenslandica*; *Ta*, *Trichoplax adhaerens*; *Nv*, *Nematostella vectensis*; *Hm*, *Hofstenia miamia*; *Dm*, *Drosophila melanogaster*; *Sp*, *Strongylocentrotus purpuratus*; *Lv*, *Lytechinus variegatus*; *Am*, *Apis mellifera*; *Pd*, *Platynereis dumerilii*; *Hs*, *Homo sapiens*; *Mm*, *Mus musculus*; *Bf*, *Branchiostoma floridae*; *Sk*, *Saccoglossus kowalevskii*. Blue boxes highlight the critical residues for the interactions between Axin and GSK-3β.

**Supplementary Figure 6.**
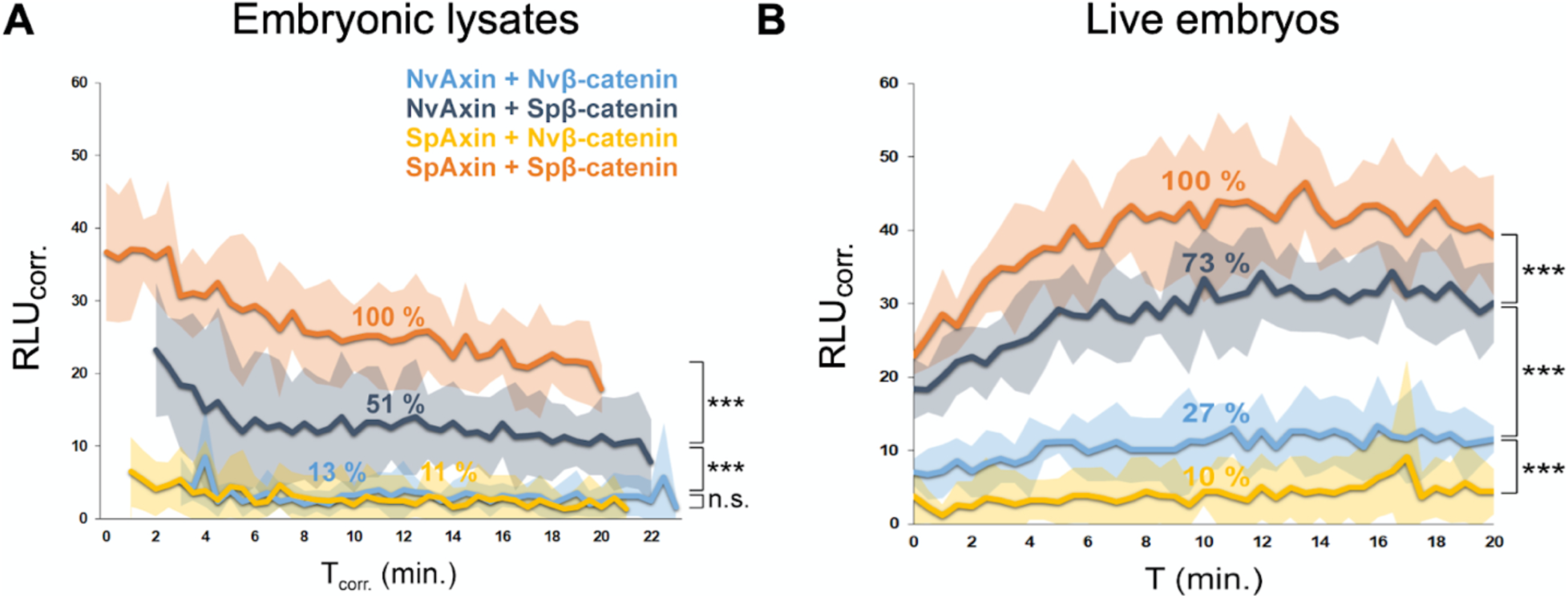
The interaction of Nv- and SpAxin with Nv- or Spβ-catenin proteins in *Nematostella* embryos or embryonic lysates. Nv/SpAxin and Nv/Spβ-catenin proteins were fused to click beetle green (CBG) split luciferase fragments, and luciferase activity was detected in embryo lysates (A) and live embryos (B) (n ≈ 150-300 per sample). Axin proteins were labeled C-terminally with the N-terminus of CBG (CBG-N) while β-catenin proteins were labeled N-terminally with the C-terminus of CBG (CBG-C), each connected by a flexible linker peptide. Proximity caused by protein-protein interaction results in complementation of CBG split-luc fragments into functional luciferase and hence stronger luminescence. Plots show moving averages (lines) and 95% confidence intervals (shaded areas) from three independent experiments that represent biological and technical replicates. Percentages are relative intensities compared to a combination of constructs with the strongest signal (SpAxin + Spβ-catenin) averaged across all replicates and timepoints. RLU_corr._ – intensity of luminescence in relative luciferase units normalized to the amount of expressed protein in each sample (as determined by Nanoluciferase signal). T, T_corr_ – time of measurement (corrected by the delay necessary to lyse embryos) *** p < 0.001, n.s. – not significant, Student’s t-test.

**Supplementary Figure 7.**
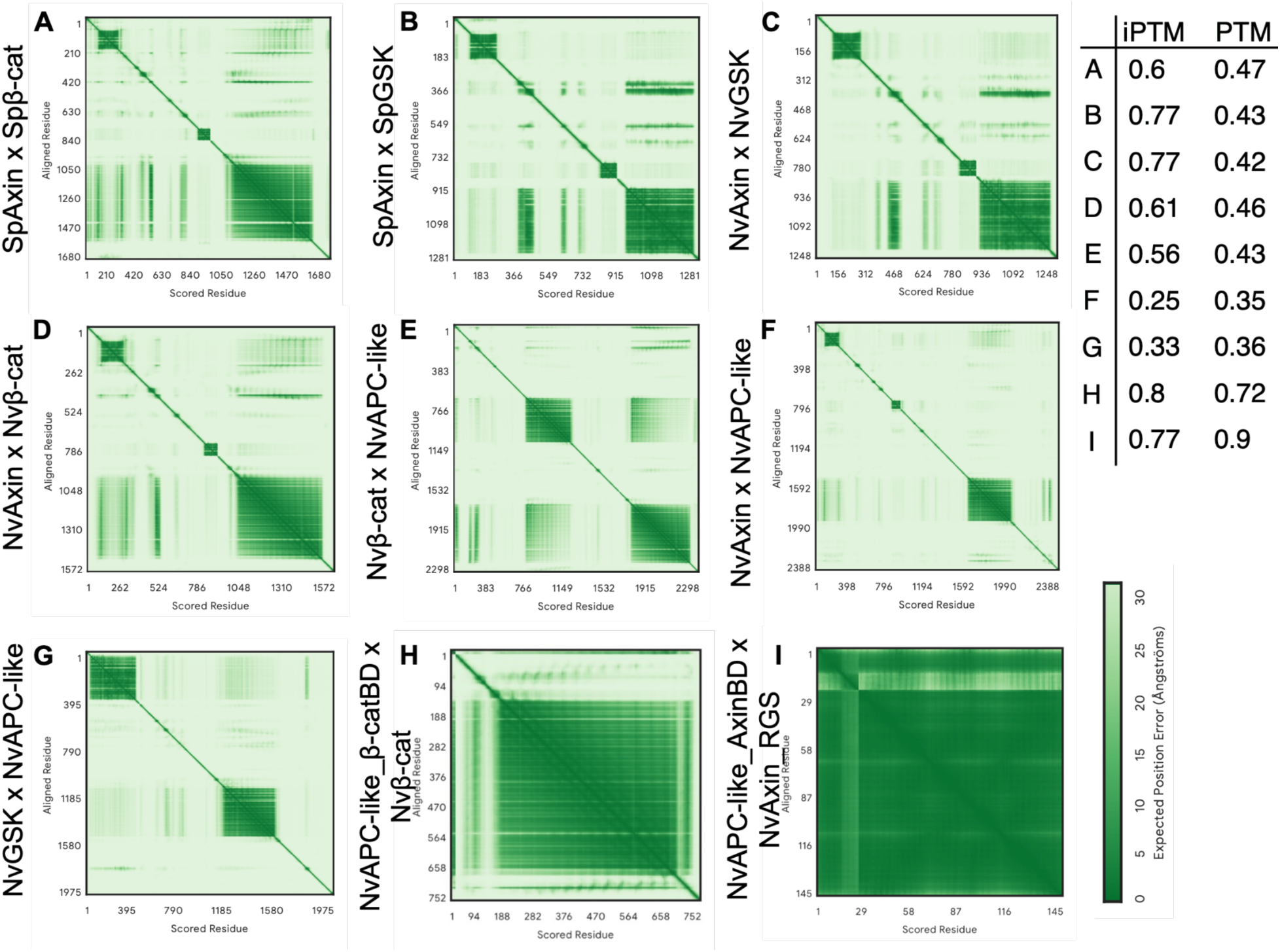
AlphaFold-based predictions of protein-protein interactions between Nvβ-catenin/Spβ-catenin and components of the β-catenin destruction complex. The expected aligned error (PAE) plots for models of interactions between (A) SpAxin and Spβ-catenin (Spβ-cat), (B) SpAxin and SpGSK, (C) NvAxin and NvGSK-3β, (D) NvAxin and Nvβ-cat, (E) Nvβ-cat and NvAPC-like, (F) NvAxin and NvAPC-like, (G) NvGSK-3β and NvAPC-like, (H) NvAPC-like_β-catBD and Nvβ-cat, (I) NvAPC-like Axin binding domain (NvAPC-like_AxinBD) and NvAxin RGS. In the PAE plots, the dark green regions indicate low position error (high confidence in the relative positioning of the interacting residues), whereas light green regions indicate higher positional error (lower confidence). The iPTM and PTM scores, which represent confidence in the predicted inter-protein and intra-protein interactions, are also provided for each model on the right side of figure.

**Supplementary Figure 8.**
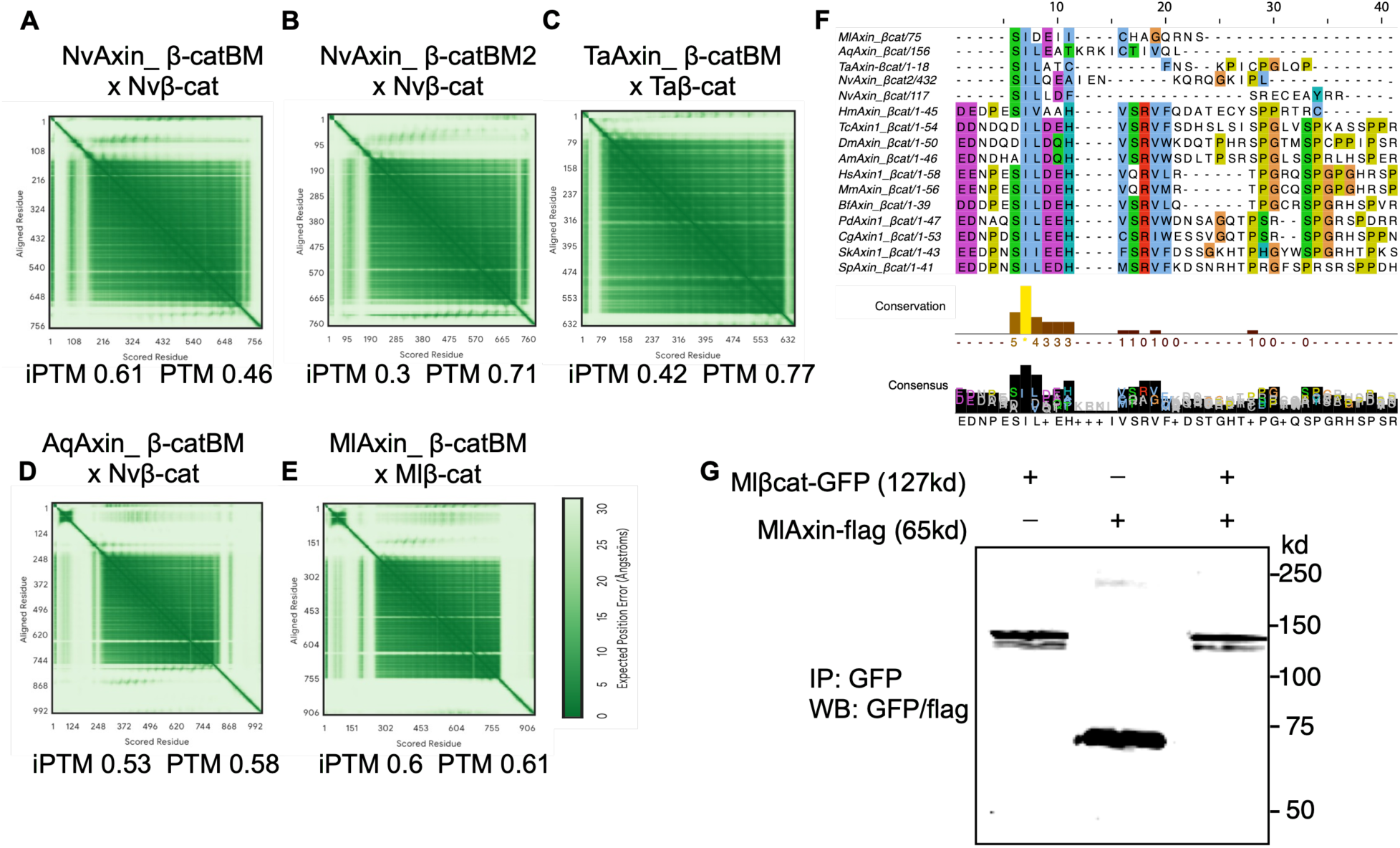
Prediction of non-bilaterian Axin-β-catenin binding motif (βcatBM) using AlphaFold 3. (A-E) The Predicted Aligned Error (PAE) plots show the predicted interactions between NvAxin_βcatBM or NvAxin_βcatBM2 (duplicated βcatBM in NvAxin) and Nvβ-catenin, TaAxin_βcatBM and Taβ-catenin, AqAxin_βcatBM and Aqβ-catenin, MlAxin_βcatBM and Mlβ-catenin. In the PAE plot, dark green regions indicate low positional error (high confidence in the relative positioning of the interacting residues), while light green regions indicate higher positional error (lower confidence). The iPTM and PTM scores, which represent confidence in the predicted inter-protein and intra-protein interactions, are also provided for each model. (F) Protein sequence alignments of various bilaterian Axin-βcatBM domains with the predicted NvAxin-βcat, TaAxin-βcat, AqAxin-βcat and MlAxin-βcat motifs. Each residue in the alignment is color-coded, while gaps are shown in white. Conservation and consensus analysis are presented at the bottom of the sequence alignment. Sequence alignments were performed using Clustal Omega and Jalview. (G) An in vitro analysis of *Mnemiopsis* Axin and *Mnemiopsis* β-catenin interaction. Co-IP analysis of MlAxin-Flag and Mlβ-catenin-GFP fail to show an interaction between these two proteins in vitro (lane 3). *Ml*, *Mnemiopsis leidyi*; *Aq*, *Amphimedon queenslandica*; *Ta*, *Trichoplax adhaerens*; *Nv*, *Nematostella vectensis*; *Hm*, *Hofstenia miamia*; *Tc*, *Tribolium castaneum*; *Dm*, *Drosophila melanogaster*; *Am*, *Apis mellifera*; *Hs*, *Homo sapiens*; *Mm*, *Mus musculus*; *Bf*, *Branchiostoma floridae*; *Pd*, *Platynereis dumerilii*; *Cg*, *Crassostrea gigas*; *Sk*, *Saccoglossus kowalevskii*; *Sp*, *Strongylocentrotus purpuratus*.

**Supplementary Figure 9.**
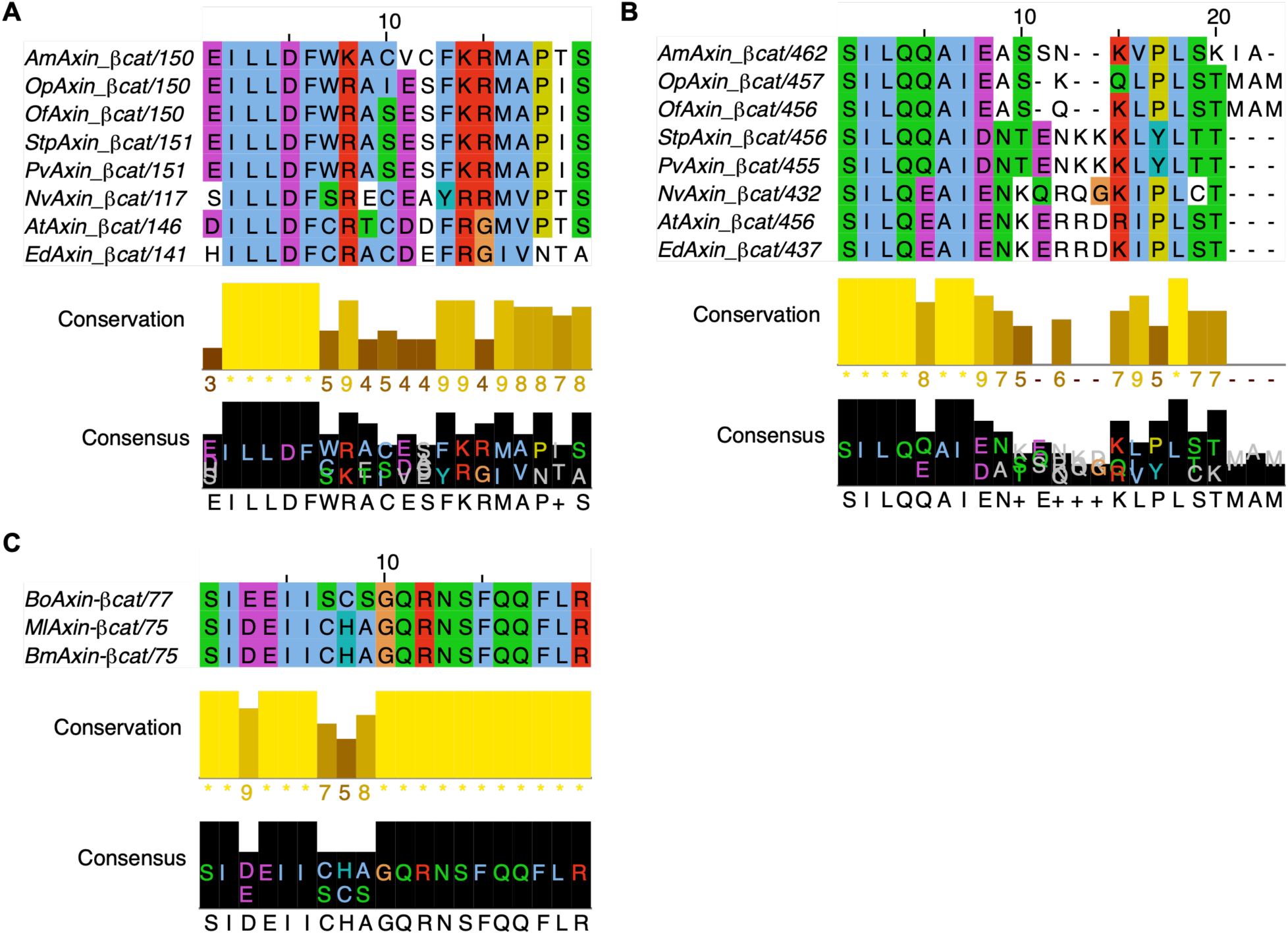
The protein sequence alignment of Axin-β-catenin binding motif (βcatBM) -like sequence in cnidarians and ctenophores. The top panels show the alignments of amino acids sequence of βcatBM-like in Axin-RGS (A) or C-terminal (B) generated using Clustal Omega and visualized with Jalview in cnidarian Axins. (C) Similar alignment of the βcatBM-like in Axin-RGS from three ctenophore species. Conserved amino acids are highlighted in distinct colors, while gaps are shown in white. The bottom panels in A, B and C show the analysis of conservation and consensus derived from the alignment. *Nv, Nematostella vectensis; Ed, Exaiptasia diaphana; At, Actinia tenebrosa; Op, Oculina patagonica; Am, Acropora muricata; Stp, Stylophora pistillata; Of, Orbicella faveolata; Pv, Pocillopora verrucosa; Ml, Mnemiopsis leidyi; Bo, Beroe ovata; Bm, Bolinopsis microptera*.

